# The genome sequence of *Aloe vera* reveals adaptive evolution of drought tolerance mechanisms

**DOI:** 10.1101/2020.05.29.122895

**Authors:** Shubham K. Jaiswal, Abhisek Chakraborty, Shruti Mahajan, Sudhir Kumar, Vineet K. Sharma

## Abstract

*Aloe vera* is a species from Asphodelaceae plant family having unique characteristics such as drought resistance and also possesses numerous medicinal properties. However, the genetic basis of these phenotypes is yet unknown, primarily due to the unavailability of its genome sequence. In this study, we report the first *Aloe vera* draft genome sequence comprising of 13.83 Gbp and harboring 86,177 coding genes. It is also the first genome from the Asphodelaceae plant family and is the largest angiosperm genome sequenced and assembled till date. Further, we report the first genome-wide phylogeny of monocots with *Aloe vera* using 1,440 one-to-one orthologs that resolves the genome-wide phylogenetic position of *Aloe vera* with respect to the other monocots. The comprehensive comparative analysis of *Aloe vera* genome with the other available high-quality monocot genomes revealed adaptive evolution in several genes of the drought stress response, CAM pathway, and circadian rhythm in *Aloe vera*. Further, genes involved in DNA damage response, a key pathway in several biotic and abiotic stress response mechanisms, were found to be positively selected. This provides the genetic basis of the evolution of drought stress tolerance capabilities of *Aloe vera*. This also substantiates the previously suggested notion that the evolution of unique characters in this species is perhaps due to selection and adaptive evolution rather than the phylogenetic divergence or isolation.

## INTRODUCTION

*Aloe vera* is a succulent and drought-resistant plant belonging to the genus *Aloe* of family Asphodelaceae [1]. More than 400 species are known in genus *Aloe*, of which four have medicinal properties with *Aloe vera* being the most potent species [2]. *Aloe vera* is a perennial tropical plant with succulent and elongated leaves having a transparent mucilaginous tissue consisting of parenchyma cells in the center referred to as *Aloe vera* gel [3]. The plant is extensively used as a herb in traditional practices in several countries, and in cosmetics and skin care products due to its pharmacological properties including anti-inflammatory, anti-tumor, anti-viral, anti-ulcers, fungicidal, etc. [4, 5]. These medicinal properties emanate from the presence of numerous chemical constituents such as anthraquinones, vitamins, minerals, enzymes, sterols, amino acids, salicylic acids, and carbohydrates [6, 7]. These properties make it commercially important, with a global market worth 1.6 billion [8].

One of the key characteristics of this succulent plant is drought resistance that enables it to survive in adverse hot and dry climates [1]. The plant has thick leaves arranged in an attractive rosette pattern to the stem. As an adaptation to the hot climate, the plant is able to perform a photosynthetic pathway known as crassulacean acid metabolism (CAM) that helps in limiting the water loss by transpiration [9]. Moreover, the leaves have the capacity to store a large volume of water in their tissues [10]. It is also known to synthesize more of soluble carbohydrates to make the osmotic adjustments under the limited water conditions, thus improving the water use efficiency [11]. Though several studies have been performed on drought stress tolerance and potential benefits of *Aloe vera*, the unavailability of its reference genome sequence has been a deterrent in understanding the genetic basis and molecular mechanisms of the unique characteristics of this medicinal plant.

In addition to the functional analysis, the resolution of the phylogenetic position has the potential to reveal the evolutionary history, and to understand the correlations between phylogenetic diversity and important traits of interest. Multiple attempts have been made to resolve the phylogenetic position of *Aloe* genus and *Aloe vera*, however, these efforts only used a few conserved loci such as rbcL, psbA, matK, and ribosomal genes, and could not be performed at genome-wide level due to the unavailability of the genomic sequence [2, 12, 13]. The previous phylogenies have reported that *Aloe vera* shared the most common recent ancestor with the species of Poales and Zingiberales order, also within the Asparagales order, it was closest to the other succulent genera such as *Haworthia*, *Gasteria*, and *Astroloba* [14, 15].

The unavailability of the genome sequence of *Aloe vera* is also noteworthy given the fact that the representative genomes of species from almost all the plant families, including Brassicaceae, Cannabaceae, Cucurbitaceae, Euphorbiaceae, Fabaceae, Malvaceae, Rosaceae, Solanaceae, Poaceae, Orchidaceae, Betulaceae have been sequenced and studied. However, till date, none of the species from the Asphodelaceae plant family has been sequenced. However, an estimate of the genome size of *Aloe vera* is available in the Plant DNA c-value database, estimated as 16.04 Gbp with a diploid ploidy level containing 14 (2n) chromosomes [16]. Thus, the availability of *Aloe vera* genome sequence will help to reveal the genomic signatures of Asphodelaceae family and will also be useful in understanding the genetic basis of the important phenotypes such as medicinal properties and drought resistance in *Aloe vera*.

Therefore in this study, we report the first draft genome sequence of *Aloe vera* using a hybrid sequencing and assembly approach by combining the Illumina short-read and oxford nanopore long-reads sequences to construct the genome sequence. The transcriptome sequencing and analysis of two tissues, root and leaf, was carried out to gain deep insights into the gene expression and to precisely determine its gene set. The genome-wide phylogeny of *Aloe vera* with other available monocot genomes was also constructed to resolve its phylogenetic position. The comparative analysis of *Aloe vera* with other monocot genomes revealed adaptive evolution in its genes and provided insights on the stress tolerance capabilities of this species.

## METHODS AND MATERIALS

### Sample collection and sequencing

The *Aloe vera* plant was bought from a plant nursery in Bhopal, India. The pulp or gel from the leaf was scrapped out and the rest was used for the DNA extraction followed by amplification of complete ITS1 and ITS2 (Internal Transcribed Spacer) and Maturase K (MatK) regions for species identification. The library was prepared using NEBNext Ultra II DNA Library preparation Kit for Illumina (New England Biolabs, England) and TruSeq DNA Nano Library preparation kits (Illumina, Inc., United States). The libraries were sequenced on Illumina HiSeq X ten and NovaSeq 6000 platforms (Illumina, Inc., United States) to generate 150 bp paired-end reads. The DNA extraction for long read sequencing was performed as per the Oxford nanopore protocols. The purified samples were used for library preparation by following the protocol of Genomic DNA by Ligation using SQK-LSK109 kit (Oxford Nanopore, UK). The library was loaded on FLO-MIN106 Flow cell (R 9.4.1) and sequenced on MinION (Oxford Nanopore, UK) using MinKNOW software (versions 3.4.5 and 3.6.0). The leaf and root part of plant were used for RNA extraction using TRIzol reagent (Invitrogen, USA). The library was prepared by using TruSeq Stranded mRNA LT Sample Prep kit and following TruSeq Stranded mRNA Sample Preparation Guide (Illumina, Inc., United States) and sequenced on Illumina NovaSeq 6000 platform for 101 bp paired end reads. Prior to sequencing the quality and quantity of libraries were assessed using Agilent 2100 Bioanalyzer (Agilent Technologies, Santa Clara, CA) and qPCR, respectively. The detailed methodology and protocols are mentioned in **Supplementary Text S1**.

### Genome assembly

The raw Illumina sequence data was processed using the Trimmomatic V0.38 tool [17]. For nanopore data, the adapter sequences were removed by using Porechop v0.2.3. SGA-preqc was used to estimate the genome size of *Aloe vera* using a k-mer count distribution method [18]. The filtered paired and unpaired Illumina reads were *de novo* assembled using ABySS v2.1.5 [19]. Different assemblies were generated on a sample dataset at increasing k-mer values, which showed the best assembly at k-mer value of 107, and hence the final assembly on complete data was performed at this k-mer value. The preprocessed nanopore reads were *de novo* assembled using wtdbg2 v2.0.0 [20]. The obtained genome assembly was first corrected for the assembly and sequencing errors using short-read data by SeqBug [21]. The hybrid assembly from the short-read and long-read assembly was generated by considering only those contigs from the ABySS and wtdbg2 assemblies that showed less than 50% query coverage and 90% identity using BLASTN against each other. The RNA-seq data based scaffolding was performed using ‘Rascaf’, followed by the long-read based gap-closing performed using LR_Gapcloser to generate the final *Aloe vera* genome assembly [22]. The other details about the data preprocessing, genome size estimation, and genome assembly and polishing are mentioned in **Supplementary Text S2 and Supplementary Figure S1**.

### Genome annotation

The genome annotation was performed on all the contigs of hybrid assembly. The tandem repeats were identified using Tandem Repeat Finder (TRF) v4.09 [23]. The microRNAs (miRNAs) were identified using a homology-based approach using miRBase database, and tRNAs were predicted using tRNAscan-SE v2.0.5 [24–27] (**Supplementary Text S3**).

### Transcriptome assembly

The transcriptome assembly of *Aloe vera* was carried out using the RNA-seq data generated from the root and leaf tissue in this study and previous studies [8, 28]. All the quality-filtered paired and unpaired transcriptome sequencing reads were *de novo* assembled using Trinity v2.6.6 software with default parameters to generate the assembled transcripts [29]. The transcriptome assembly was evaluated by mapping the filtered RNA-seq data on the assembled transcripts using hisat2 v2.1.0 [30]. The BUSCO score was used to assess the completeness of the transcriptome assembly calculated by BUSCO v4.0.5 software using the standard database specific to the Liliopsida class [31, 32].

### Gene set construction

The maker pipeline was used for gene set construction of the *Aloe vera* genome [33]. The soft-masked genome of *Aloe vera* (contigs ≥300 bp) generated using RepeatMasker v4.1.0 with Repbase repeat library (RepeatMasker Open-4.0, http://www.repeatmasker.org) was used for the gene prediction using the maker pipeline. Both the *ab initio* and empirical evidence were used for the gene predictions. The *Aloe vera* EST evidence from the RNA-seq assembly of *Aloe vera* species, protein sequences of the closest species *Dioscorea rotundata* and *Musa acuminata*, and *ab initio* gene predictions of the *Aloe vera* genome were used to construct the gene set using the maker pipeline. AUGUSTUS v3.2.3 was used for the *ab initio* gene prediction, and the BLAST alignment tool was used for homology-based gene prediction using the EST evidence in the maker pipeline [34–36]. Further, Exonerate v2.2.0 was used to polish and curate the BLAST alignment results (https://github.com/nathanweeks/exonerate). The evidence from *ab initio* and homology-based methods were integrated to perform the final gene predictions.

The genes from predicted transcripts were identified by extracting the longest isoforms. The unigenes were identified by performing the clustering using CD-HIT-EST v4.8.1 program, and coding regions were predicted using TransDecoder v5.5.0 (https://github.com/TransDecoder/TransDecoder) [37–41]. The gene set constructed using the maker pipeline and transcriptome assembly was filtered, and only the genes with ≥300 bp length were considered further. The clustering of remaining maker pipeline based genes was performed using CD-HIT-EST v4.8.1 program with 95% identity and a seed size of 8 bp [41]. The transcriptome gene set was searched in the maker gene set using BLASTN. The genes from the transcriptome assembly gene set that matched to the maker gene set with the parameters: identity ≥50%, e-value <10^−9^, and query coverage ≥50% were removed. The remaining genes for the transcriptome assembly gene set were directly added to the maker gene set to construct the final gene set of *Aloe vera*.

### Orthogroups identification

For orthogroups identification, the representative of monocot species from all the clades, for which high-quality genomes were available on Ensembl plants database, were selected along with an outgroup species, the model plant *Arabidopsis thaliana*. The selected monocot species were *Aegilops tauschii*, *Brachypodium distachyon*, *Dioscorea rotundata*, *Eragrostis tef*, *Hordeum vulgare*, *Leersia perrieri*, *Musa acuminata*, *Oryza sativa*, *Panicum hallii fil2*, *Saccharum spontaneum*, *Setaria italica*, *Sorghum bicolor*, *Triticum aestivum*, and *Zea mays*. The proteome files containing all the protein sequences of the 15 species retrieved from Ensembl plants release 46 [42], and the protein-coding genes from the transcriptome assembly of *Aloe vera* were used to construct the orthogroups. The longest transcript for each gene was extracted for each species using in-house python scripts. The proteome files with longest transcripts were used for the orthogroups identification using OrthoFinder v2.3.9 [43]. The OrthoFinder v2.3.9 analysis included a total of 16 species, i.e., 14 monocot species, the model species *Arabidopsis thaliana* as an outgroup, and *Aloe vera* sequenced in this study.

### Orthologous gene set construction

From the orthogroups identified by the OrthoFinder analysis, the orthogroups with the taxon count of 16 were extracted, which included genes from each of the 16 species. A total of 5,472 orthogroups were extracted using this criterion. Only the longest gene of each species was retained in each of these orthogroups to construct the orthologous gene set. Thus, a total of 5,472 orthologs were identified across 16 species. From these 5,472 orthologs one-to-one orthologs were extracted. To include maximum number of genes in the one-to-one orthology, the fuzzy one-to-one orthogroups instead of true one-to-one orthogroups were identified using KinFin v1.0 [44]. A total of 1,440 one-to-one orthologs were extracted using this method across the selected 16 species.

### Phylogenetic tree construction

The phylogenetic species tree was constructed with the fuzzy one-to-one orthologous genes. The individual orthologous sets were aligned using MAFFT v7.455 [45]. The alignments were trimmed using BeforePhylo v0.9.0 (https://github.com/qiyunzhu/BeforePhylo) to remove the poorly aligned regions. All protein sequence alignments of orthologs across 16 species were concatenated using BeforePhylo v0.9.0, followed by species phylogenetic tree construction using RAxML v8.2.12 [46]. The maximum likelihood phylogenetic tree was constructed using the rapid hill climbing algorithm with the 100 bootstrap replicates. Since the amino acid sequences were used, the ‘PROTGAMMAGTR’ substitution model was utilized to construct the species tree.

### Identification of genes with a higher rate of evolution

The genes that show higher root-to-tip branch length are considered to have a higher rate of nucleotide divergence or mutation, indicating a higher rate of evolution. For this analysis, the individual maximum likelihood phylogenetic trees were constructed using the protein sequences of the 5,472 orthologs identified across the 16 species. The maximum likelihood phylogenetic trees with 100 bootstrap replicates were constructed using the rapid hill climbing algorithm with the ‘PROTGAMMAGTR’ substitution model by using RAxML v8.2.12 [46]. The root-to-tip branch length values were calculated for each of the 16 extant species using the ‘adephylo’ package in R [47, 48]. All the genes that showed a significantly higher root-to-tip branch length for *Aloe vera* in comparison to rest of the species were extracted using in-house Perl scripts and were considered to be the genes with a higher rate of evolution in *Aloe vera*.

### Identification of positively selected genes

The positively selected genes in *Aloe vera* were identified using the branch-site model implemented in the PAML software package v4.9a [49]. An iterative program for sequence alignment, SAT’e, was utilized to perform the alignments of the 5,472 ortholog protein sequences. The combination of Prank, MUSCLE, and RaxML was used to perform the SAT’e based alignment to control the false positives and false negatives in the alignment [50]. The protein-alignment guided codon alignment was performed for the 5,472 ortholog nucleotide sequences using ‘TRANALIGN’ program of EMBOSS v6.5.7 package [51]. The ‘codeml’ was run on ortholog codon alignments using the species phylogenetic tree constructed in previous steps. The alignments were filtered for the ambiguous codon sites and gaps and only the clean sites were considered for the positive selection analysis. The likelihood ratio tests were performed using the log-likelihood values for the null and alternative models, and the p-values were calculated based on the *χ*^2^-distribution. Further, the FDR corrected p-values or FDR q-values were also calculated. All genes with FDR-corrected p-values <0.05 were considered to be the genes with positive selection in *Aloe vera*. Further, all codon sites with >0.95 probability of being positively selected in the ‘foreground’ branch based on the Bayes Empirical Bayes analysis were considered to be the positively selected codon sites in a gene.

### Identification of genes with unique substitutions that have functional impact

The genes with unique amino acid substitutions in *Aloe vera* species in comparison to all the selected species were identified. The protein alignments for the 5,472 orthologs were generated using the MAFFT v7.455 [45]. The positions that are identical in all the species but different in *Aloe vera* were identified and considered to be the sites with unique amino acid substitutions in *Aloe vera*. In this analysis, the gaps were ignored, and also the sites with gaps present in the 10 amino acids flanking regions on both sides were ignored. This step helped in considering only the sites with proper alignment for the unique substitution analysis. The identification of unique amino acid sites was performed by using the in-house python scripts. The functional impact of the unique amino acid substitutions on the protein function was identified using the Sorting Intolerant From Tolerant (SIFT) tool with UniProt database as reference [52, 53].

### Identification of genes with multiple signs of adaptive evolution (MSA)

The genes that showed at least two signs of adaptive evolution among the three signs of adaptive evolution tested above (higher rate of evolution, positive selection, and unique substitution with functional impact) were considered as the genes with multiple signs of adaptive evolution or MSA genes in *Aloe vera*.

### Functional annotation

The functional annotation of gene sets was performed using multiple methods. The functional annotation and functional categorization of genes into different eggNOG categories was performed using the eggNOG-mapper [54]. The genes were assigned to different KEGG pathways, and also the KEGG orthology was determined using the most updated KAAS genome annotation server [55]. The gene ontology enrichment or GO term enrichment analysis was performed using the WebGestalt web server [56]. In the over representation analysis, only the GO categories with the p-value <0.05 in the hypergeometric test were considered to be functionally enriched in the gene set. Further, the functional annotation of genes was also manually curated. The assignment of genes to the specific categories and phenotypes was performed by manual annotation. The protein-protein interaction and co-expression data were extracted from the STRING database, and the network analysis was performed using Cytoscape [57, 58].

## RESULTS

### Sequencing of *Aloe vera* genome and transcriptome

The estimated genome size of *Aloe vera* is 16.04 Gbp, and to comprehensively cover this large genome, a total of 506.4 Gbp (~32X) of short-reads and 123.5 Gbp (~7.7X) of long-reads data was generated using Illumina and nanopore platforms, respectively (**Supplementary Table S1 and S2**) [16, 59]. For transcriptome, a total of 6.6 Gbp and 7.3 Gbp of RNA-seq data was generated from leaf and root, respectively. The transcriptome data from this study and the publicly available RNA-seq data from previous studies [8, 28] were combined together, resulting in a total of 37.1 Gbp of RNA-seq data for *Aloe vera*, which was used for the analysis (**Supplementary Table S3**). All the genomic and RNA-seq read data were trimmed and filtered using Trimmomatic, and only the high-quality read data was used to construct the final genome and transcriptome assemblies. The complete workflow of the sequence analysis is shown in **Supplementary Figure S1**.

### Assembly of *Aloe vera* genome

The final draft genome assembly of *Aloe vera* had the size of 13.83 Gbp with N50 and largest scaffold of 3.18 kbp and 4.94 Mbp, respectively (**Supplementary Table S4**). Of which, 12.25 Gbp had length >300bp with N50 of 7.03 kbp, and 9.85 Gbp had length >500bp with N50 of 13.06 kbp, which is a challenging feat for such a gigantic plant genome, and is also comparable to the other large plant genomes assembled till date [60–63]. This was achieved by the hybrid assembly of short-read and long-read data, which was further polished by correction using SeqBug, RNA-seq based scaffolding using Rascaf, and long-read based gapclosing using LR-gapcloser. The k-mer count distribution-based method using only the short Illumina reads estimated a genome size of 13.63 Gb, which was smaller than the c-value-based genome size estimation of 16.04 Gbp, conceivably due to the usage of only short reads data for the genome size estimation (**Supplementary Figure S2**). The %GC for the final assembly was 41.98%. The analysis of repetitive sequences revealed 557,638,058 bp of tandem repeats corresponding to 3.41% of the complete genome.

### Transcriptome assembly

The Trinity assembly of transcriptomic reads resulted in a total size of 163,190,792 bp with an N50 value of 1,268 bp and an average contig length of 796 bp (**Supplementary Table S5**). The mapping of filtered RNA-seq reads on the Trinity transcripts using hisat2 resulted in the overall percentage mapping of 92.49%. The complete BUSCO score (addition of single copy and duplicates) on the transcripts was 87.7%. A total of 205,029 transcripts were predicted, corresponding to 108,133 genes with the percent GC of 43.69. The clustering of gene sequences using CD-HIT-EST to remove the redundancy resulted in 107,672 unigenes. The coding genes (CDS) from the unigenes were predicted using TransDecoder resulting in 34,269 coding genes.

### Genome annotation and gene set construction

A total of 1,978 standard amino acid specific tRNAs and 378 hairpin miRNAs were identified in the *Aloe vera* genome (**Supplementary Table S6**). The maker pipeline-based gene prediction resulted in a total of 114,971 coding transcripts, of which 63,408 transcripts (≥300 bp) were considered further for clustering at 95% identity resulting in 57,449 unique coding gene transcripts. Application of the same length-based selection criteria (≥300 bp) on trinity-identified 34,269 coding gene transcripts resulted in 33,998 coding gene transcripts. The merging to these two coding gene transcript sets resulted in the final gene set of 86,177 genes for *Aloe vera*, which had the complete BUSCO score of 69.0% and single copy BUSCO score of 65.7%.

### Identification of orthologous across selected plant species

A total of 104,543 orthogroups were identified using OrthoFinder across the selected 16 plant species, of which 9,343 orthogroups were unique to *Aloe vera* and contained genes only from *Aloe vera*. Only a total of 5,472 orthogroups had sequences from all the 16 plant species and were used for the identification of orthologs. For these 5,472 orthogroups, in case of presence of more than one gene from a species in an orthogroup, the longest gene representative from that species was selected to construct the final orthologous gene set for any orthogroups. Thus, including one gene from each of the 16 species in an orthogroup, a total of 5472 orthologs were identified. In addition, the fuzzy one-to-one orthologs finding approach applied using KinFin resulted in a total of 1,440 fuzzy one-to-one orthologs that were used for constructing the maximum likelihood species phylogenetic tree.

### Resolving the phylogenetic position of *Aloe vera*

Each of the 1,440 fuzzy one-to-one orthologous gene set was aligned and concatenated, and the resulted concatenated alignment had a total of 1,453,617 alignment positions. The concatenated alignment was filtered for the undetermined values, which were treated as missing values, and a total of 1,157,550 alignment positions were retained. The complete alignment data and the filtered alignment data were both used to construct maximum likelihood species trees using RAxML with the bootstrap value of 100, and both the alignment data resulted in the same phylogeny. Thus, the phylogeny based on the filtered data was considered to be the final genome-wide phylogeny of *Aloe vera* with all the representative monocot genomes available on Ensembl plants database and *Arabidopsis thaliana* as an outgroup (**Figure 1**). This phylogeny also corroborated with the earlier reported phylogenies by Silvera et al., 2014, Dunemann et al., 2014, and Wang et al., 2016, which were constructed using a limited number of genetic loci [64–66]. It is apparent from the phylogeny that *Dioscorea rotundata* and *Musa acuminata* are the most closely related to *Aloe vera*, and share the same clade (**Figure 1**). All other selected monocots are distributed in separate clade with *Triticum aestivum* and *Aegilops tauschii* being the most distantly related to *Aloe vera*.

**Figure 1.**
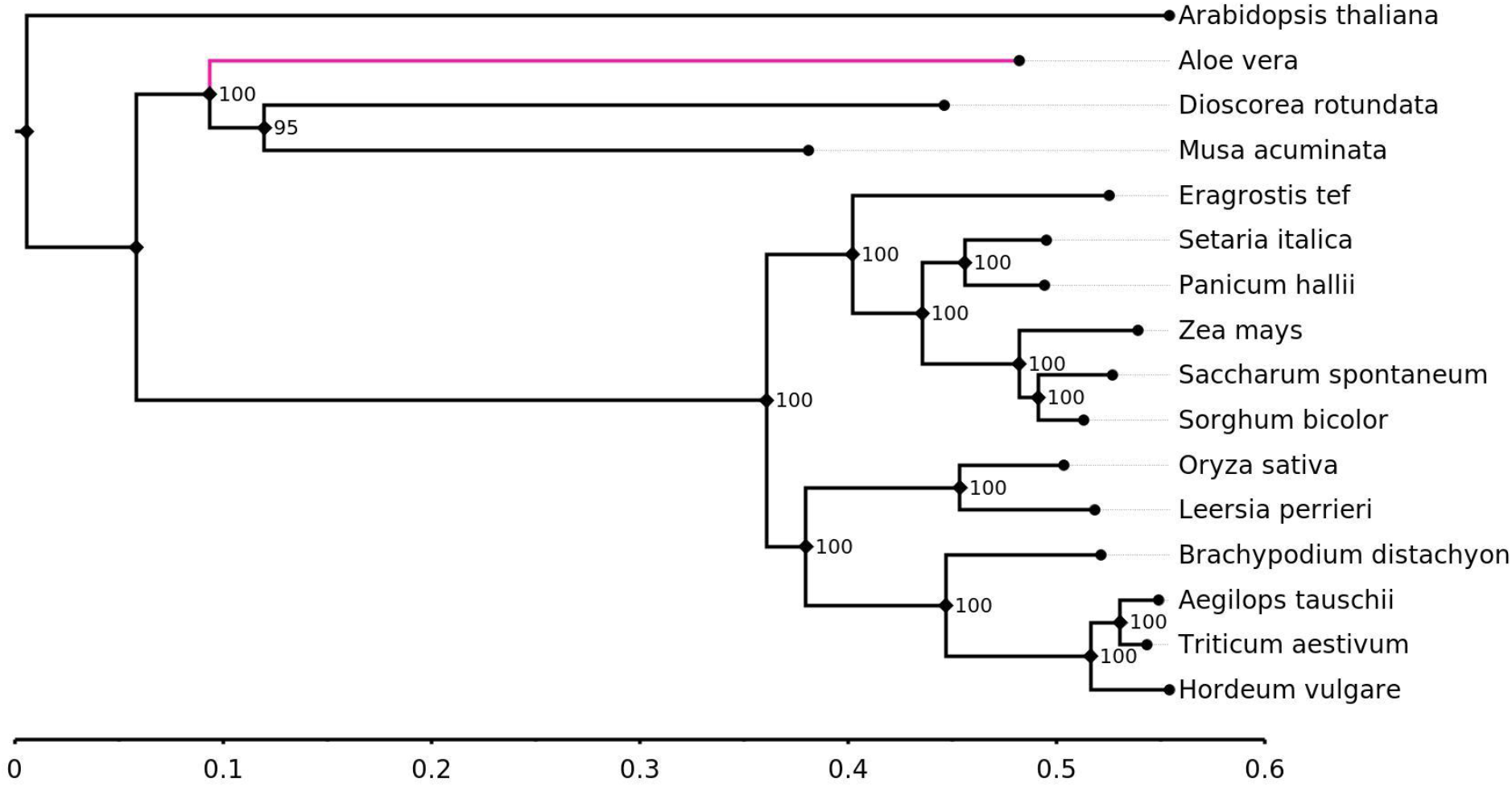
The phylogenetic tree of the selected 14 monocot species, *Aloe vera*, and *Arabidopsis thaliana* as an outgroup The values mentioned at the nodes are the bootstrap values. The scale mentioned is the nucleotide substitutions per base.

Recently an updated plant megaphylogeny has been reported for the vascular plants [14]. The species of Poales order showed similar relative positions in our reported phylogeny and this megaphylogeny. In the megaphylogeny, *Musa acuminata* was reported to share the most common recent ancestor with the species of Poales order, but in our phylogeny we observed that *Musa acuminata* shared the most common recent ancestor with *Dioscorea rotundata* from Dioscoreales order (**Figure 1 and Supplementary Figure S3**). Also, among the selected monocots, the species of Dioscoreales order was reported to show the earliest divergence. However, in our genome-wide phylogeny, *Aloe vera* showed the earliest divergence.

Also, with respect to the reported phylogeny of angiosperms, at the order level the Poales and Zingiberales formed a clade, and their ancestor shared the most recent common ancestor with Asparagales, then all three shared a recent ancestor with Dioscoreales [15]. In our genome-wide phylogeny, Zingiberales and Dioscoreales shared the most recent common ancestor, and their ancestor shares the most recent common ancestor with Asparagales, and the three shared a recent ancestor with Poales.

### Genes with a higher rate of evolution

A total of 85 genes showed higher rates of evolution in *Aloe vera* in comparison to the other monocot species. These genes belonged to several eggNOG categories and KEGG pathways, as mentioned in **Supplementary Table S7 and Supplementary Table S8,** with a higher representation of ribosomal genes. The distribution of enriched (p-value<0.05) biological process GO terms is mentioned in **Supplementary Table S9**. Also, among these 85 genes, three molecular function GO terms, rRNA binding, structural constituent of cytoskeleton, and structural constituent of ribosome showed an enrichment (p-value<0.05) (**Supplementary Table S10**). Five transcription factors WRKY, MYB, bHLH, CPP, and LBD showed higher rates of evolution in *Aloe vera*. Among these, WRKY, MYB, and bHLH are known to be involved in drought stress tolerance [67–69]. There were six chloroplast functioning related genes, namely EMB3127, PnsB3, TL29, IRT3, PDV2, and SIRB, that showed a higher rate of evolution. Notably, the chloroplast function related genes have been implicated in different abiotic stress conditions in plants, including drought [70, 71].

### Identification of positively selected genes

A total of 199 genes showed positive selection in *Aloe vera* with the FDR q-value threshold of 0.05. The distribution of these genes in eggNOG categories, KEGG pathways, and GO term categories are mentioned in **Supplementary Table S11-S15**. Among the genes with positive selection, several genes were involved in key functions with specific phenotypic consequences (**Figure 2**). These included flowering related genes that are important for the reproductive success, calcium-ion binding and transcription factors/sequence-specific DNA binding genes involved in signal transduction for response to external stimulus, carbohydrate catabolism genes required for energy production, and genes involved in abiotic stress response [72–74]. Among the abiotic stress response genes, there were four categories of genes that were involved in response to water-related stress, DNA damage response genes involved in reactive oxidative species (ROS) stress response, nuclear pore complex genes involved in plant stress response by regulating the nucleo-cytoplasmic trafficking, and secondary metabolites biosynthesis related genes that deal with different types of biotic and abiotic stresses [75–77]. The robust and efficient DNA damage response mechanism is essential for biotic and abiotic stress tolerance, and for the genomic stability [78]. Thus, adaptive evolution in this pathway seemingly contributes towards the stress tolerance capabilities and genomic stability in *Aloe vera*.

**Figure 2.**
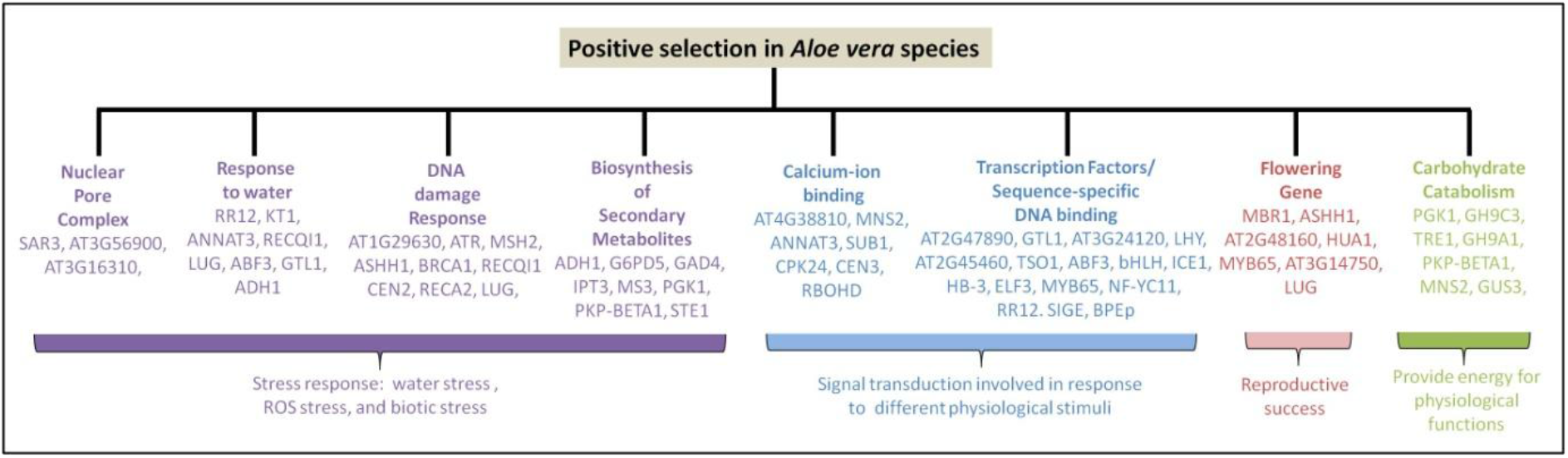
The functional categories of genes that showed positive selection in *Aloe vera* The standard *Arabidopsis thaliana* gene IDs were used in case of genes that did not have a standard gene symbol.

Another gene G6PD5 that showed positive selection in *Aloe vera* protects plants against different types of stress, such as salinity stress by producing nitric oxide (NO) molecule, which leads to the expression of Defence response genes [79, 80]. Regulation of osmotic potential under drought stress is acquired by different ion channels, transporters, and carrier proteins [81]. In this study, K^+^ transporter 1(KT1), bidirectional amino acid transporter 1(BAT1), and Sodium Bile acid symporter (AT4G22840) genes were found to be positively selected in *Aloe vera*.

The Abscisic acid (ABA) responsive element binding factor (ABF) gene was found to be positively selected. This gene is differentially expressed under drought and other abiotic stress and alters specific target gene expression by binding to ABRE (abscisic acid-response element), the characteristic element of ABA-inducible genes [82]. ABA also regulates stomatal closure and solute transport, and thus have implications in drought tolerance [83]. The trehalase 1 (TRE1) gene was also found to be positively selected, and the over-expression of this gene causes better drought tolerance through ABA guided stomatal closure [84].

### Genes with site-specific signs of evolution

Two types of site-specific signatures of adaptive evolution i.e., positively selected codon sites and unique amino acid substitutions with significant functional impact were identified in *Aloe vera*. A total of 1,848 genes had positively selected codon sites, and a total of 2,669 genes had unique amino acid substitutions with functional impact. The distribution of genes with positively selected codon sites and unique amino acid substitutions with functional impact in eggNOG categories, KEGG pathways, and GO term categories are mentioned in **Supplementary Tables S16-S25**.

One of the characteristics of succulent plants such as *Aloe vera* is the ability to efficiently assimilate the atmospheric CO_2_ and reduce water loss by transpiration through the crassulacean acid metabolism (CAM) pathway a specific mode of photosynthesis. The evolution of CAM is an adaptation to the limited CO_2_ and limited water condition, and a significant correlation between higher succulence and increased magnitude of CAM metabolism has been observed [85]. In this study, several crucial genes of CAM metabolism showed site-specific signatures of adaptive evolution in *Aloe vera* (**Figure 3**). The potassium channel involved in stomatal opening/closure (KAT2), malic enzyme (ME) that converts malic acid to pyruvate, and phosphoenolpyruvate carboxylase (PEPC) that converts phosphoenolpyruvate to oxaloacetate and assimilates the environmental CO_2_ showed both the signs of site-specific adaptive evolution. In addition, the other CAM genes including potassium transport 2/3 (KT2/3), pyruvate orthophosphate dikinase (PPDK), phosphoenolpyruvate carboxylase kinase 1 (PPCK1), carbonic anhydrase 1 (CA1), peroxisomal NAD-malate dehydrogenase 2 (PMDH2), tonoplast dicarboxylate transporter (TDT), and aluminum activated malate transporter family protein (ALMT9) showed unique substitutions with functional impact in *Aloe vera*.

**Figure 3.**
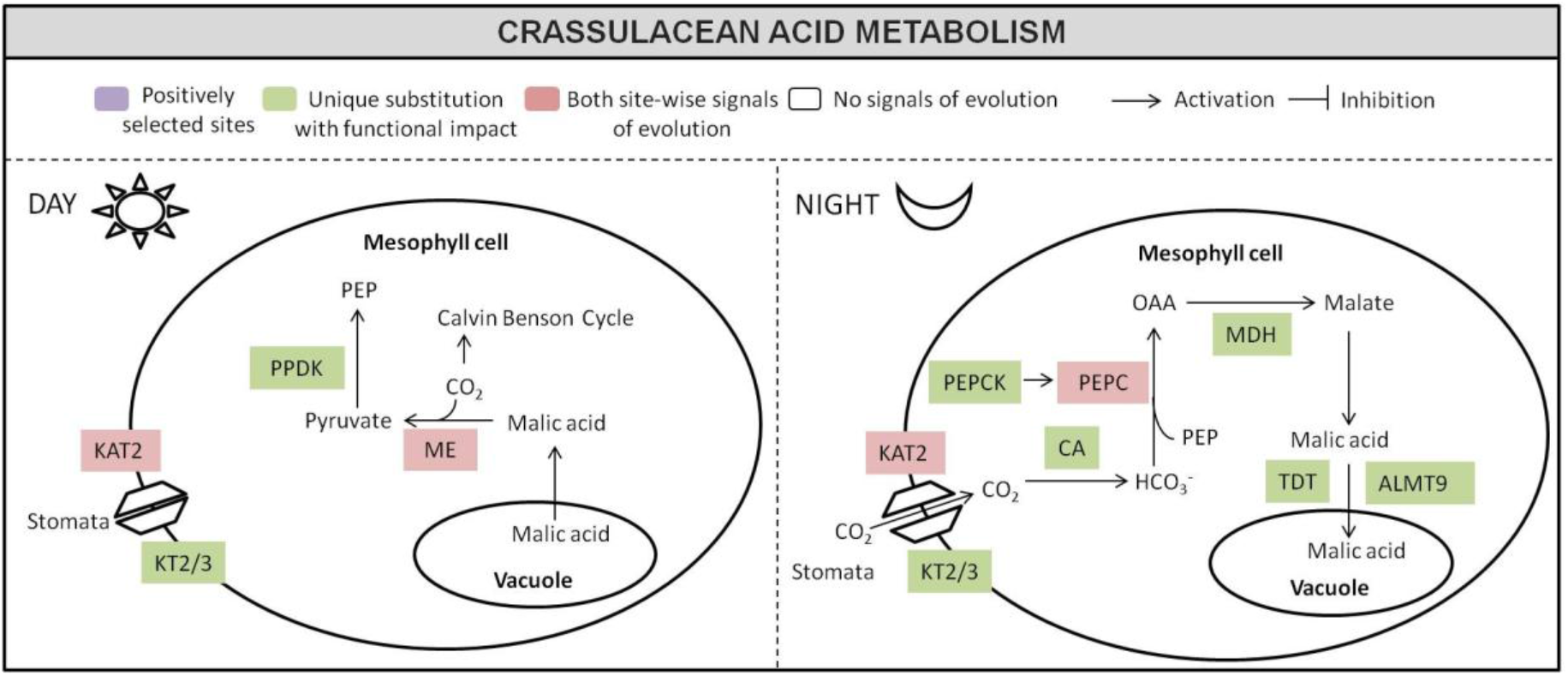
The adaptive evolution of CAM pathway in *Aloe vera* The important genes of the CAM pathway are shown with their function in the day time and night time metabolism. The genes in Levander color had positively selected codon sites, the genes in Green color had unique substitutions with function impact, and the genes in Red color showed both the signs of site-specific adaptive evolution in *Aloe vera*. There were no CAM pathway genes that had only positively selected codon sites.

CAM metabolism evolution is known to be a result of modified circadian regulation at the transcription and posttranscriptional levels [86]. CAM evolution is the well-characterized physiological rhythm in plants, and it is also a specific example of circadian clock-based specialization [86, 87]. Several plant circadian rhythm genes showed site-specific signs of adaptive evolution in *Aloe vera* (**Figure 4**). Three essential genes of red light response, PHYB, ELF3, and LHY, showed both the signs of site-specific adaptive evolution. Also, the FT gene important for flowering and under the control of circadian rhythm showed both the signs of site-specific adaptive evolution. The PHYA gene, which is also a part of the red light response, had unique substitutions with functional impact. Among the blue light response genes, three genes GI, FKF1, and SPA2 had unique substitutions with functional impact, and two genes HY5 and CHS had positively selected codon acid sites. The blue light response regulates the UV-protection and photomorphogenesis.

**Figure 4.**
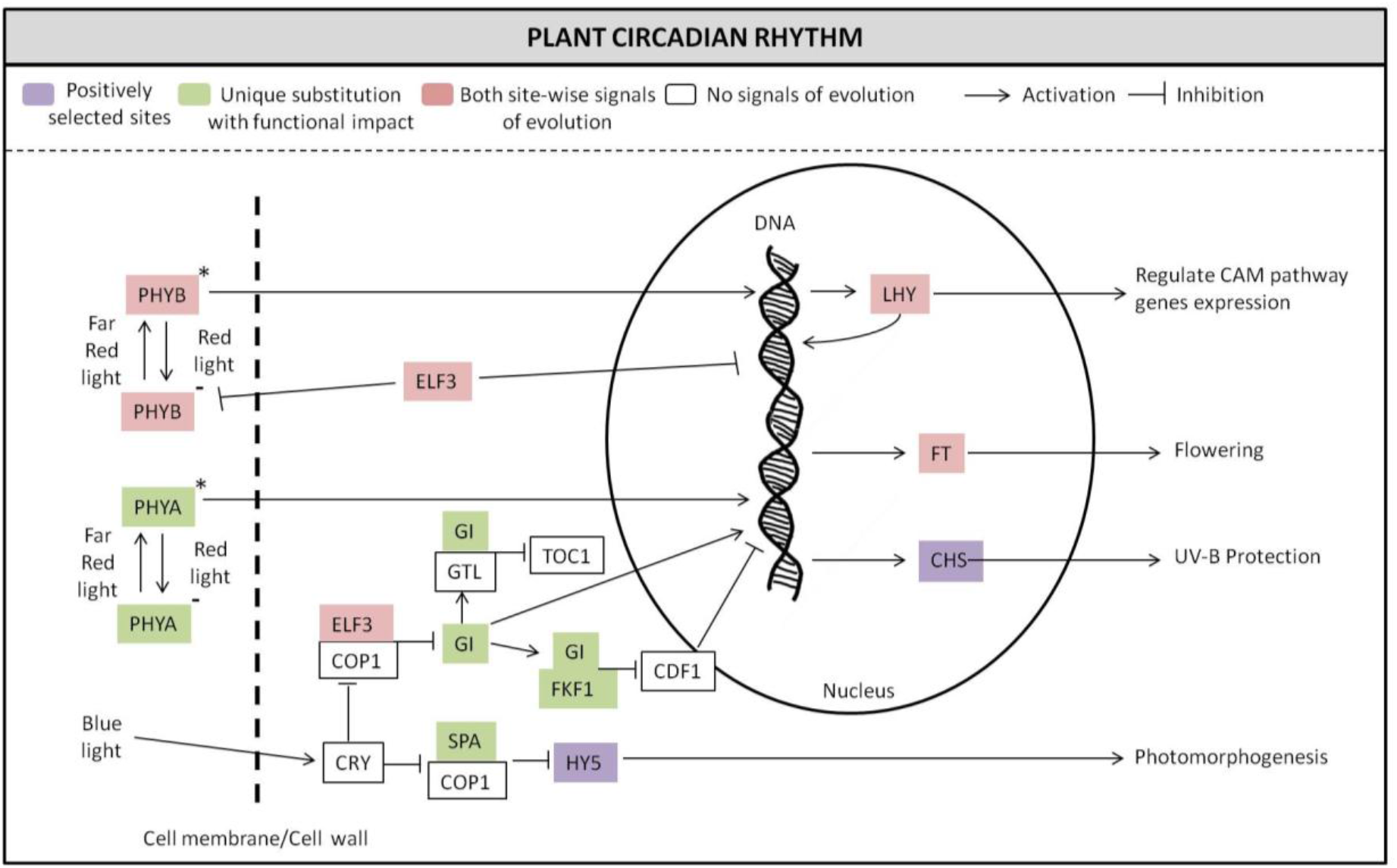
The adaptive evolution of circadian rhythm pathway in *Aloe vera* The important genes of the plant circadian rhythm are shown with their function. The genes in Levander color had positively selected codon sites, the genes in Green color had unique substitutions with function impact, and the genes in Red color showed both the signs of site-specific adaptive evolution in *Aloe vera*.

Plant hormone signaling regulates plant growth, development, and response to different types of biotic and abiotic stress [88]. Multiple genes of auxin, cytokinin, and brassinosteroid hormone signaling involved in cellular growth and elongation having implications in cellular and tissue succulence, showed site-specific signatures of adaptive evolution (**Figure 5**). The genes of the abscisic acid hormone signaling involved in stomatal opening/closure required for CAM metabolism and different biotic and abiotic stress response [82] had positively selected amino acid sites and unique substitution sites with functional impact (**Figure 5**). Also, the genes involved in salicylic acid signaling important for providing disease resistance and help in biotic stress response showed site-specific signatures of adaptive evolution (**Figure 5**).

**Figure 5.**
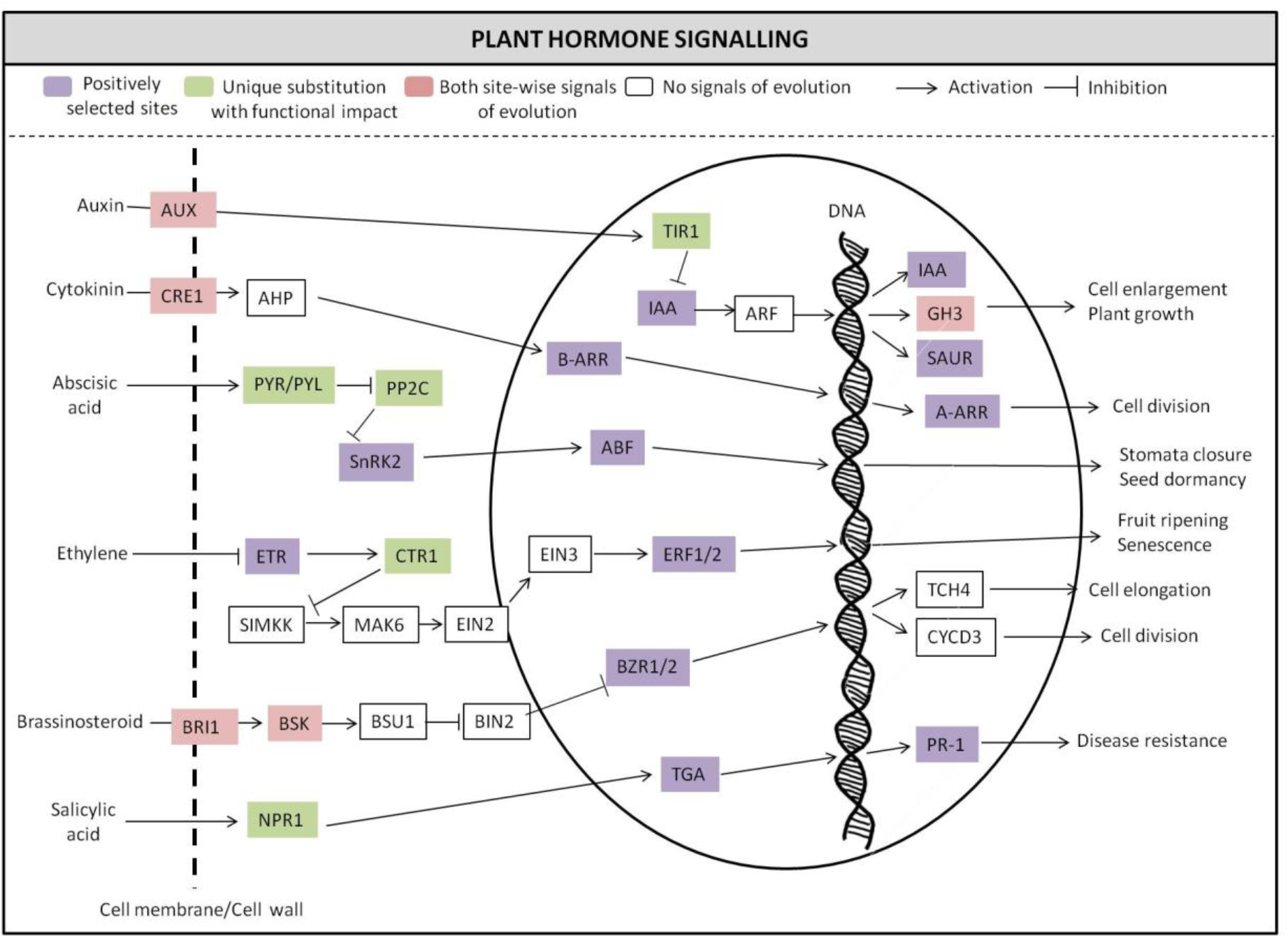
The adaptive evolution of plant hormone signaling pathway in *Aloe vera* The important genes of the auxin, cytokinin, abscisic acid, ethylene, brassinosteroid, and salicylic acid signaling pathways are shown with their function. The genes in Levander color had positively selected codon sites, the genes in Green color had unique substitutions with function impact, and the genes in Red color showed both the signs of site-specific adaptive evolution in *Aloe vera*.

### Genes with multiple signs of adaptive evolution

Among the three signatures of adaptive evolution i.e., positive selection, a higher rate of evolution, and unique amino acid substitutions with functional impact, a total of 148 genes showed two or more signs of adaptive evolution and were identified as the genes with multiple signs of adaptive evolution (MSA). The distribution of these genes in eggNOG categories, KEGG pathways, and GO categories are mentioned in **Supplementary Table S26-S29**. Another study that performed the proteomic analysis of drought stress response in wild peach also found similar categories to be enriched in the proteins that were differentially expressed under drought conditions [89]. A total of 112 genes out of the 148 MSA genes in *Aloe vera* were from the specific categories that are involved in providing drought stress tolerance. The specific groups of proteins and their relation with the drought stress tolerance are mentioned in **Figure 6**.

**Figure 6.**
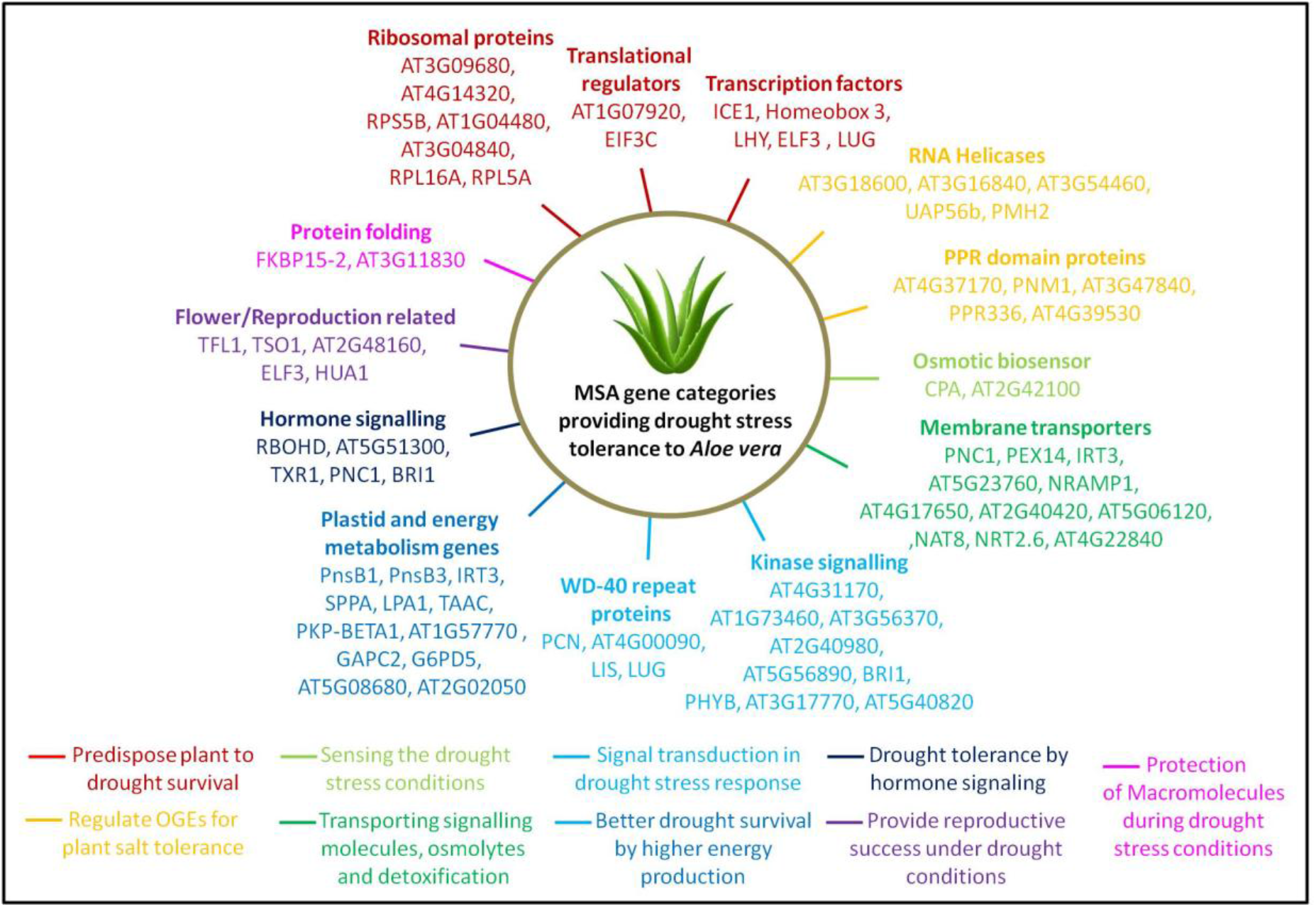
The MSA genes in *Aloe vera* that are involved in drought stress response The relation of specific categories of genes with drought stress response was determined from the literature. The standard *Arabidopsis thaliana* gene IDs were used in case of genes that did not have a standard gene symbol.

Several ribosomal genes, translational regulators, and transcription factors genes were found to be MSA genes in this study, and these were also found to be over-expressed under drought conditions in different proteomic and transcriptomic studies and aid in better drought stress survival [89, 90]. Many nuclear genes are involved in the functioning of symbiotic organelles chloroplast and mitochondria. Among these genes, some genes are also involved in the organellar gene expression (OGEs) regulation, and mutants of these genes are known to show altered response to different abiotic stress, including high salinity stress [91, 92]. Several of these genes belonging to two categories, RNA helicases and PPR domain proteins, were found to be MSA genes. Thus, in the *Aloe vera* species, these genes have been adaptively evolved to provide this species with better salt tolerance.

Two osmotic biosensor genes, ‘CPA’ and ‘AT2G42100’, were found to be among the MSA genes in *Aloe vera*. Different membrane transporters that can transport signaling molecules, osmolytes, and metals were also among the MSA genes (**Figure 6**). These included two peroxisomal transporters ‘PNC1’ a nucleotide carrier protein, and ‘PEX14’ a transporter for PTS1 and PTS2 domain containing signaling proteins, different heavy metal transporters such as ‘IRT3’ an iron transporter, ‘AT5G23760’ a copper transporter, and ‘NRAMP1’ a manganese transporter, and ‘AT4G17650’ a lipid transporter, ‘AT2G40420’ an amino acid transporter, ‘AT5G06120’ an intracellular protein transporter, ‘ALA1’ a phospholipid transporter, ‘NAT8’ a Nucleobase-ascorbate transporter, ‘NRT2.6’ a high-affinity nitrate transporter, and ‘BASS6’ a sodium/metabolite co-transporter. These osmotic sensors and transporters provide significant enhancement in function in drought stress condition and help in adjusting to the water scarcity [93, 94].

The genes for several kinases and WD-40 repeat proteins were also found to be among the MSA genes in *Aloe vera*. These proteins are involved in signaling and transcription regulation required for the drought stress tolerance [90, 95–97]. Also, the genes involved in energy generation and are part of the thylakoid membrane showed MSA. The stability of thylakoid membrane proteins has been associated with drought resistance, and these energy production related genes are crucial in survival during the drought stress [90, 98]. Two genes that assist in protein folding were found to show MSA (**Figure 6**), and these proteins are very important in protecting the macromolecules of the cells under the drought stress conditions [99]. Five genes involved in plant hormone signaling were also among the MSA genes. The plant hormone signaling is central to the signaling pathways required for the drought stress tolerance [100]. Five genes involved in flowering and reproduction regulation were also found to be among the MSA genes in *Aloe vera*. The flowering and reproduction related genes are known to be regulated for better reproductive success under drought stress conditions as part of the drought tolerance strategy used by many plants [101, 102].

The co-expression of MSA genes was examined using the co-expression data from the STRING database [57], and the MSA genes that co-express with at least one other MSA gene are displayed as a network diagram (**Figure 7A**). From the network, it is evident that almost all co-expressing MSA genes are drought stress tolerance related, and the genes forming the dense network are also drought stress tolerance related. Predominantly, three categories of drought stress tolerance related MSA genes have shown co-expression: genes involved in energy production, genes involved in OGEs regulation, and genes that predispose plants to drought stress tolerance.

**Figure 7.**
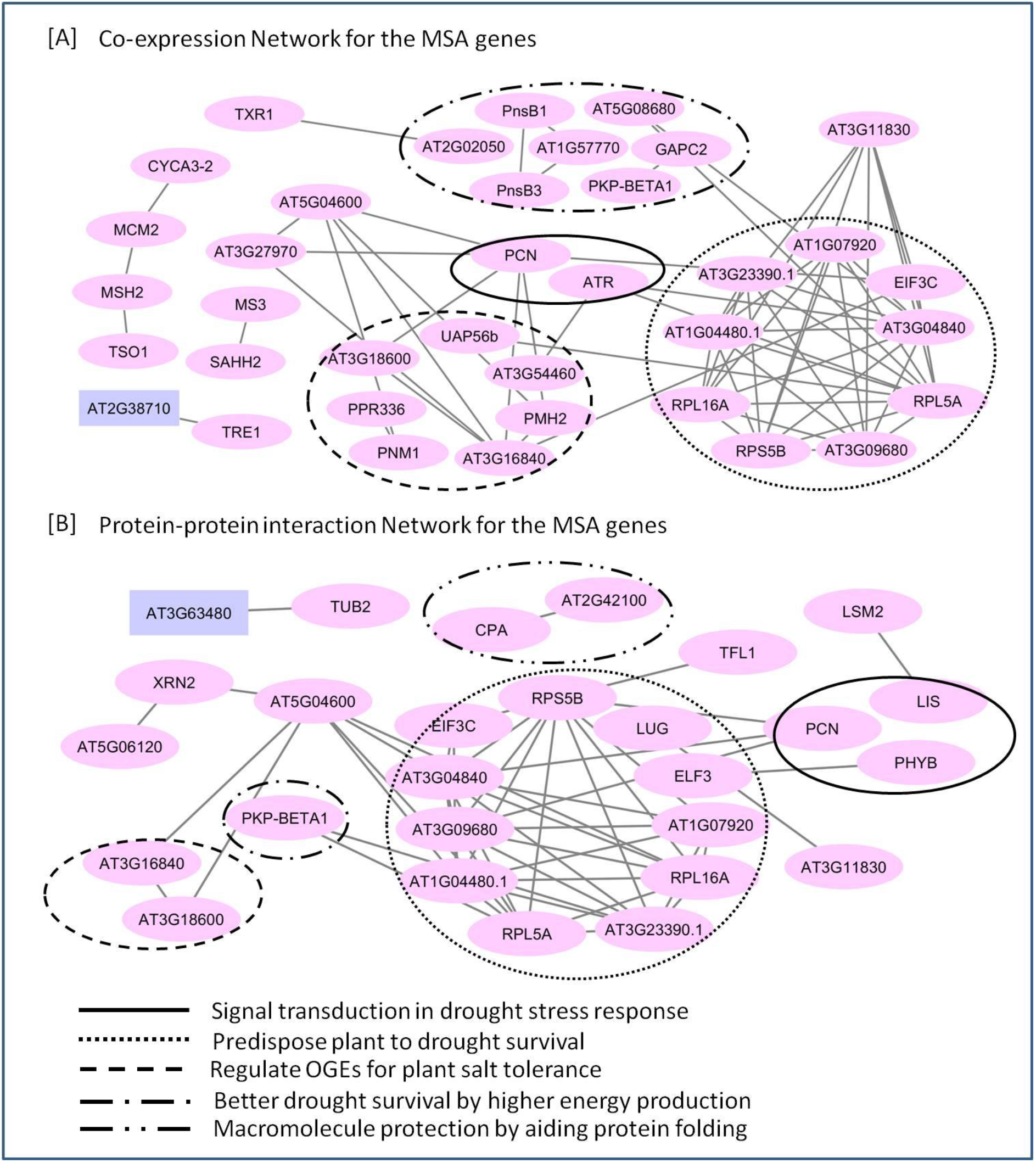
Evaluating the co-expression and physical interaction of MSA genes in *Aloe vera* [A] The co-expression network of the MSA genes is shown. Only the MSA genes that showed at least one co-expression connection are shown. The nodes represent the genes, and the edges represent the co-expression of the connected nodes. [B] The protein-protein interaction network of the MSA genes is shown. Only the MSA genes that showed at least one protein-protein interaction are shown. The nodes represent the genes, and the edges represent the protein-protein interaction between the connected nodes. Note: The standard *Arabidopsis thaliana* gene IDs were used in case of genes that did not have a standard gene symbol.

Similarly, a network diagram was constructed using the protein-protein interaction data of MSA genes from the STRING database [57]. The genes with physical interaction known from the experimental studies are shown in **Figure 7B**. From the network, it is apparent that among the interacting MSA genes, all of them except one are involved in drought stress tolerance. Further, among the MSA genes involved in drought stress tolerance, it is primarily the genes that predispose plants to drought stress tolerance, and the genes involved in signal transduction in drought stress response showed the physical interaction. In addition, two genes that function as osmotic biosensors and two OGEs regulation genes also displayed physical interaction.

## DISCUSSION

In this work, we have presented the complete draft genome sequence of *Aloe vera*, which is an evolutionarily important, ornamental, and widely used plant species due to its medicinal properties, pharmacological applications, traditional usage, and commercial value. The availability of *Aloe vera* genome sequence is also important since it is the first genome sequenced from the Asphodelaceae plant family, and is the largest angiosperm and the fifth largest genome sequenced so far. It is also the largest genome sequenced using the oxford nanopore technology till date. The hybrid approach of using short-read (Illumina) and long-read (nanopore) sequence data emerged as a successful strategy to tackle the challenge of sequencing one of the largest plant genomes.

The study reported the gene set of *Aloe vera* constructed using the combination of *de novo* and homology-based gene predictions, and also using the data from the genomic assembly and the transcriptomic assembly from multiple tissues, thus indicating the comprehensiveness of the approach. The *Aloe vera* had a higher number of coding genes than the other monocots used in this study except for *Triticum aestivum*, which had more number of coding genes (**Supplementary Table S30**). The estimation of coding genes in *Aloe vera* was similar to the number of genes in other monocot genomes suggesting the correctness of the gene prediction and estimation.

This study reported the first genome-wide phylogeny of *Aloe vera* with all other monocot species available on the Ensembl plant database, and with *Arabidopsis thaliana* as an outgroup. A few previous studies have also examined the phylogenetic position of *Aloe vera* with respect to other monocots but used a few genomic loci. Thus, this is the first genome-wide phylogeny of monocots that resolves the phylogenetic position of *Aloe vera* with respect to the other monocots by using 1,440 different loci distributed throughout their genomes. The very high bootstrap values for the internal nodes and existence of no polytomy in the phylogeny further attest to the correctness of the phylogeny. This phylogeny is mostly in agreement with the previously known phylogenies, and also provided some new insights [2, 14, 64–66].

An earlier phylogeny constructed using “ppc-aL1a” gene showed that *Sorghum bicolor, Zea mays, Setaria italica, Brachypodium distachyon, Hordeum vulgare*, and *Oryza sativa* form a monophyletic group, which was also observed in our phylogeny [66]. Similarly, the relative positions of *Hordeum vulgare, Saccharum officinarum, Zea mays*, and *Oryza sativa* in another phylogeny based on “CENH3” gene were in agreement with our phylogeny [64]. Using the “NORK” gene, another recent study reported the relative phylogenetic position of four monocot species: *Oryza sativa, Zea mays, Sorghum bicolour*, and *Setaria italica* [65]. *Zea mays, Sorghum bicolour*, and *Setaria italica* were found to share a recent last common ancestor and *Oryza sativa* had diverged earlier from their common ancestor, which is also supported by the genome-wide phylogeny reported in this study.

Though the genome-wide phylogeny showed the species of Poales order with similar topology as reported in earlier studies, a different topology was observed for the relative position of Musa acuminata, *Dioscorea rotundata*, and *Aloe vera* from the orders Zingiberales, Dioscoreales, and Asparagales, respectively [14, 15]. The observed differences could be due to the usage of a few genomic loci in the previous phylogenies, whereas the phylogeny reported in this study is a genome-wide phylogeny constructed using 1,440 one-to-one orthologs distributed across the genome. The availability of more complete genomes from monocots and the inclusion of more genomic loci in the phylogenetic analysis will help explain the observed differences and confirm the relative positions of these species.

One of the key highlights of the study was the revelation of adaptive evolution of genes involved in drought stress response, which provides a genetic explanation for the drought stress tolerance properties of *Aloe vera*. This plant is known to display a number of phenotypes such as perennial succulent leaves and CAM mechanism for carbon fixation that provide it with better drought stress survival [10]. Several experimental studies have also reported that it can make adjustments such as increased production of sugars and increased expression of heat-shock and ubiquitin proteins for efficient water utilization and osmotic maintenance that eventually provide better drought survival [11, 103, 104]. In this study, the majority (80%) of genes that showed multiple signs of evolution (MSA) were involved in drought stress tolerance related functions. These genes were also found to be co-expressing and physically interacting with each other, which further point towards the adaptive evolution of the drought stress tolerance mechanisms in this species. The adaptive evolution of genes involved in drought stress tolerance provides insights into the genetic basis of the drought resistance property of *Aloe vera*.

Further, several crucial genes of the CAM pathway and circadian rhythm have also shown site-specific signs of adaptive evolution in *Aloe vera* in comparison to the other monocot species. The CAM pathway has very high water use efficiency, and is known to have evolved convergently in many arid regions for better drought survival [105]. Also, the CAM pathway is a physiological rhythm with temporal separation of atmospheric CO_2_ assimilation and Calvin-Benson cycle, and is under the control of plant circadian rhythm [86, 106]. This CAM pathway evolution is known to be a specific type of circadian rhythm specialization [87, 107]. Thus, the observed adaptive evolution of CAM pathway and its controller circadian rhythm in this study point towards its role in providing this species an evolutionary advantage for efficient drought stress survival.

The evolutionary success of the *Aloe* genus is also known to be due to the succulent leaf Mesophyll tissue [2]. Particularly, the medicinal use of *Aloe vera* is much associated with the succulent leaf mesophyll tissue, and a loss of this tissue leads to the loss of medicinal properties [3]. The plant species with CAM pathway have large vacuoles in comparison to the non-CAM plants, and therefore the leaf succulence is also higher in CAM plants. Thus, it is tempting to speculate that the observed evolution of CAM pathway in *Aloe vera* may also be crucial for the higher leaf mesophyll succulence contributing to its medicinal properties. Also previously, it has been proposed that the specific properties of *Aloe vera* such as the high leaf succulence, medicinal properties, and drought resistance are the consequence of evolutionary processes such as selection and speciation rather than due to phylogenetic diversity or isolation [2]. The signatures of adaptive evolution in drought tolerance and CAM pathway genes in *Aloe vera* further substantiate this notion.

## CONCLUSION

The first draft genome, transcriptome, gene set, and functional analysis of *Aloe vera* reported in this study will act as a reference for future studies to understand the medicinal or evolutionary characteristics of this species, and its family Asphodelaceae. The first genome-wide phylogeny of *Aloe vera* and other available monocot genomes resolved the phylogenetic position of *Aloe vera* and emphasized the need for the availability of more genomes for precise phylogenetic analysis. The comparative genomic analyses of *Aloe vera* with the other monocot genomes provided novel insights on the adaptive evolution of drought stress response, CAM pathway, and circadian rhythm genes in *Aloe vera*, and suggest that the positive selection and adaptive evolution of specific genes contribute to the unique phenotypes of this species.

## LIST OF ABBREVIATIONS

MSA: Multiple signs of adaptive evolution
CAM: Crassulacean acid metabolism
COG: Clusters of Orthologous Groups
KEGG: Kyoto Encyclopedia of Genes and Genomes
GO: Gene ontology
BUSCO: Benchmarking Universal Single-Copy Orthologs
SIFT: Sorting Intolerant From Tolerant
FDR: False discovery rate
BLAST: Basic Local Alignment Search Tool
N50: minimum contig length needed to cover 50% of the genome
ABA: Abscisic acid
snoRNA: small nucleolar RNA
snRNA: small nuclear RNA
tRNA: transfer RNA
rRNA: ribosomal RNA
srpRNA: signal recognition particle RNA
miRNA: micro RNA
MYB: Myeloblastosis
bHLH: basic helix–loop– helix
CPP: cysteine-rich polycomb-like protein
LBD: Lateral Organ Boundaries (LOB) Domain
EMB3127: Embryo Defective 3127
PnsB3: Photosynthetic NDH subcomplex B3
TL29: Thylakoid Lumen 29
IRT3: Iron regulated transporter 3
PDV2: Plastid Division2
SIRB: Sirohydrochlorin ferrochelatase B
G6PD5: Glucose-6-phosphate dehydrogenase 5
KAT2: Potassium channel in *Arabidopsis thaliana* 2
PHYB: Phytochrome B
ELF3: Early Flowering 3
LHY: Late Elongated Hypocotyl
FT: Flowering locus T
PHYA: Phytochrome A
GI: Gigantea
FKF1: Flavin-binding, Kelch repeat, F box 1
SPA1: Suppressor of PHYA-105 1
HY5: Elongated Hypocotyl5
CHS: Chalcone synthase
CPA: Capping Protein A
PNC1: Peroxisomal adenine nucleotide carrier 1
PEX14: Peroxin 14
IRT3: Iron regulated transporter 3
NRAMP1: Natural Resistance-Associated Macrophage Protein 1
ALA1: Aminophospholipid ATPase 1
NAT8: Nucleobase-Ascorbate Transporter 8
NRT2.6: High affinity Nitrate Transporter 2.6
ppc-aL1a: Phosphoenolpyruvate carboxylase
CENH3: Centromeric histone H3
NORK: Nodulation receptor kinase
PPR: Pentatricopeptide Repeat
PTS1: Peroxisomal targeting signal 1
PTS2: Peroxisomal targeting signal 2
LTR-RT: Long terminal repeat Retrotransposons
EST: Expressed sequence tag

## AUTHORS’ CONTRIBUTIONS

VKS conceived and coordinated the project. SM prepared the DNA and RNA samples, performed sequencing, and the species identification assay. SKJ with the input from VKS designed the computational framework of the study. SKJ and AC performed the genome assembly, transcriptome assembly, genome annotation, gene set construction, orthology analysis, and species phylogenetic tree construction. SKJ performed the root-to-tip branch length, positive selection, unique substitution with functional impact, network, and statistical analysis. SKJ, AC, SK, and SM performed the functional annotation of gene sets. SKJ, AC, and VKS analysed the data. SKJ, AC, and VKS interpreted the results. SKJ and AC constructed the figures. SKJ, AC, SM, SK, and VKS wrote and revised the manuscript. All the authors have read and approved the final version of the manuscript.

## ACKNOWLEDGEMENTS

The author SKJ thanks Department of Science and Technology for the DST-INSPIRE fellowship. AC and SM thank Council of Scientific and Industrial Research (CSIR) for fellowship. SK thanks University Grants Commission (UGC) for the fellowship. We also thank the intramural research funds provided by IISER Bhopal.

