## Supplementary information for "The genome sequence of *Aloe vera* reveals adaptive evolution of drought tolerance mechanisms"

**SUPPLEMENTARY TABLES**

**Supplementary Table S1. Summary of the Illumina sequence data for *Aloe vera* genome**

| Paired-end Insert Size | Average Read Length | Number of Reads | Total Data | Sequence Coverage |
|---|---|---|---|---|
| 447 bp and 600 bp | 150 bp | 3,393,209,648 | 506.4 Gb | ~32X |

The sequencing coverage was calculated by assuming the *Aloe vera* genome size of 16.04 Gb [1, 2].

**Supplementary Table S2. Summary of the nanopore sequence data for *Aloe vera* genome**

| Average Read Length | Number of Reads | Total Data | Sequence Coverage |
|---|---|---|---|
| 2,952 bp | 41,828,185 | 123.5 Gb | ~7.7X |

The sequencing coverage was calculated by assuming the *Aloe vera* genome size of 16.04 Gb [1, 2].

**Supplementary Table S3. Summary of the transcriptome data for *Aloe vera* genome**

| Tissue | Average read length R1 (bp) | Average read length R2 (bp) | Total number of read pairs | Total number of R1 (bp) | Total number of R2 (bp) | Total number of bases (bp) |
|---|---|---|---|---|---|---|
| Leaf[1] | 101 | 101 | 32,776,695 | 3,310,446,195 | 3,310,446,195 | 6,620,892,390 |
| Root[1] | 101 | 101 | 36,212,970 | 3,657,509,970 | 3,657,509,970 | 7,315,019,940 |
| Leaf[2] | 101 | 101 | 29,247,010 | 2,953,948,010 | 2,953,948,010 | 5,907,896,020 |
| Root[2] | 145.6 | 145.4 | 51,078,070 | 7,440,880,511 | 7,427,363,139 | 14,868,243,650 |
| 1K genome project[3] | 73 | 75 | 16,218,326 | 1,183,937,798 | 1,216,374,450 | 2,400,312,248 |
| Total data | | | 165,533,071 | 18,546,722,484 | 18,565,641,764 | 37,112,364,248 |

[1]Our study, [2]Choudhri et al., 2018 [3], [3]1K project [4]

**Supplementary Table S4. Summary statistics of the final assembly for *Aloe vera* genome**

| Genome statistics* | Value |
|---|---|
| Number of scaffold (≥ 250 bp) | 15,423,352 |
| Number of scaffold (≥ 1000 bp) | 1,888,944 |
| Number of scaffold (≥ 5000 bp) | 398,067 |
| Number of scaffold (≥ 10000 bp) | 214,630 |
| Number of scaffold (≥ 25000 bp) | 76,509 |
| Number of scaffold (≥ 50000 bp) | 18,536 |
| Total length (≥ 250 bp) | 13,833,270,974 |
| Total length (≥1000 bp) | 8,950,349,048 |
| Total length (≥ 5000 bp) | 6,724,607,175 |
| Total length (≥ 10000 bp) | 5,466,494,755 |
| Total length (≥ 25000 bp) | 3,294,939,922 |

| | |
|---|---|
| Total length (≥ 50000 bp) | 1,293,064,116 |
| Largest scaffold | 4,941,938 |
| GC% (≥ 250 bp) | 41.98 |
| N50 (≥ 250 bp) | 3,180 |
| Number of N's per 100 kbp (≥ 250 bp) | 64.67 |

*On using the length criterion of ≥200 bp, the genome assembly had the size of 15.91 Gbp that covers ~99% of the genome as per the c-value-based genome size estimation of 16.04 Gbp.

**Supplementary Table S5. Summary statistics of the transcriptome assembly for *Aloe vera***

| **Statistics based on all transcript contigs** | |
|---|---|
| Contig N10 | 3151 |
| Contig N20 | 2402 |
| Contig N30 | 1942 |
| Contig N40 | 1584 |
| Contig N50 | 1268 |
| Average contig | 795.94 |
| Total assembled bases | 163,190,792 |
| **Statistics based on only longest isoform per gene** | |
| Contig N10 | 3,098 |
| Contig N20 | 2,338 |
| Contig N30 | 1,838 |
| Contig N40 | 1,431 |
| Contig N50 | 1,061 |
| Average contig | 652.68 |
| Total assembled bases | 70,576,159 |
| **Counts of genes and transcripts** | |
| Total trinity 'genes' | 108133 |
| Total trinity transcripts | 205029 |

**Supplementary Table S6. The tRNAs compared across different monocot species including *Aloe vera***

| Species | tRNA |
|---|---|
| *Aegilops tauschii* | 547 |
| *Brachypodium distachyon* | 289 |
| *Dioscorea rotundata* | 154 |
| *Hordeum vulgare* | 727 |
| *Leersia perrieri* | 196 |
| Musa acuminata | 1,254 |
| *Oryza sativa* | 242 |
| *Setaria italicaa* | 346 |
| *Sorghum bicolor* | 324 |
| *Triticum aestivum* | 1,915 |
| *Zea mays* | 2,834 |
| *Arabidopsis thaliana* | 689 |
| *Aloe vera* | 1,978* |

*Only the tRNAs specific to standard amino acids are mentioned

The data for the other species was retrieved from the Ensembl plants genome browser, and for the *Aloe vera* species the tRNAs were identified as mentioned in the **Supplementary Text S3**.

**Supplementary Table S7. The distribution of genes with higher rate of evolution in different eggNOG categories in *Aloe vera***

| eggNOG category | Number of genes |
|---|---|
| Function unknown | 17 |
| Translation, ribosomal structure and biogenesis | 9 |
| Posttranslational modification, protein turnover, chaperones | 8 |
| Inorganic ion transport and metabolism | 6 |
| Intracellular trafficking, secretion, and vesicular transport | 5 |
| Transcription | 5 |
| Energy production and conversion | 5 |
| RNA processing and modification | 5 |
| Carbohydrate transport and metabolism | 4 |
| Cytoskeleton | 4 |
| Signal transduction mechanisms | 3 |
| Coenzyme transport and metabolism | 2 |
| Amino acid transport and metabolism | 2 |
| Replication, recombination and repair | 2 |
| Nucleotide transport and metabolism | 1 |
| Lipid transport and metabolism | 1 |
| Cell cycle control, cell division, chromosome partitioning | 1 |

**Supplementary Table S8. The distribution of genes with higher rate of evolution in different KEGG pathways in *Aloe vera* (Only pathways with more than ten genes are mentioned)**

| KEGG Pathway | Number of genes |
|---|---|
| Ribosome | 8 |
| Alzheimer disease | 3 |
| Huntington disease | 3 |
| Oxidative phosphorylation | 2 |
| Glutathione metabolism | 2 |
| Spliceosome | 2 |
| RNA transport | 2 |
| Thermogenesis | 2 |
| Parkinson disease | 2 |
| Pathogenic Escherichia coli infection | 2 |
| Salmonella infection | 2 |

**Supplementary Table S9. The biological process GO categories that were enriched in the genes with higher rate of evolution in *Aloe vera* (Only statistically significant GO terms p<0.05 are mentioned)**

| GO Term ID | Description | p-value |
|---|---|---|
| GO:0051187 | cofactor catabolic process | 0.0156 |
| GO:0072511 | divalent inorganic cation transport | 0.0234 |
| GO:0009624 | response to nematode | 0.0278 |
| GO:0048285 | organelle fission | 0.0299 |
| GO:0070925 | organelle assembly | 0.0350 |
| GO:0007017 | microtubule-based process | 0.0401 |

**Supplementary Table S10. The molecular function GO categories that were found enriched in the genes with higher rate of evolution in *Aloe vera* (Only statistically significant GO terms p<0.05 are mentioned)**

| GO Term ID | Description | p-value |
|---|---|---|
| GO:0019843 | rRNA binding | 0.0014 |
| GO:0005200 | structural constituent of cytoskeleton | 0.0042 |
| GO:0003735 | structural constituent of ribosome | 0.0384 |

**Supplementary Table S11. The distribution of genes with positive selection in different eggNOG categories in *Aloe vera***

| eggNOG category | Number of genes |
|---|---|
| Function unknown | 47 |
| Transcription | 19 |
| Carbohydrate transport and metabolism | 15 |
| RNA processing and modification | 13 |
| Signal transduction mechanisms | 12 |
| Posttranslational modification, protein turnover, chaperones | 11 |
| Inorganic ion transport and metabolism | 9 |
| Intracellular trafficking, secretion, and vesicular transport | 9 |
| Translation, ribosomal structure and biogenesis | 8 |
| Replication, recombination and repair | 8 |
| Energy production and conversion | 6 |
| Amino acid transport and metabolism | 5 |
| Lipid transport and metabolism | 5 |
| Coenzyme transport and metabolism | 5 |
| Cell cycle control, cell division, chromosome partitioning | 5 |
| Secondary metabolites biosynthesis, transport and catabolism | 4 |
| Chromatin structure and dynamics | 3 |
| Cell wall/membrane/envelope biogenesis | 3 |
| Defence mechanisms | 2 |
| Cytoskeleton | 2 |
| Nuclear structure | 1 |

**Supplementary Table S12. The distribution of genes with positive selection in different KEGG pathways in *Aloe vera* (Only pathways with more than one gene are mentioned)**

| KEGG Pathway | Number of Genes |
|---|---|
| Cell cycle | 5 |
| RNA transport | 4 |
| Glycolysis / Gluconeogenesis | 3 |
| Starch and sucrose metabolism | 3 |
| alpha-Linolenic acid metabolism | 3 |
| Cysteine and methionine metabolism | 3 |
| Aminoacyl-tRNA biosynthesis | 3 |
| Plant hormone signal transduction | 3 |
| Cellular senescence | 3 |
| Circadian rhythm - plant | 3 |
| Thermogenesis | 3 |
| Plant-pathogen interaction | 3 |
| Central carbon metabolism in cancer | 3 |
| Human papillomavirus infection | 3 |

| | |
|---|---|
| Fructose and mannose metabolism | 2 |
| Pyruvate metabolism | 2 |
| Photosynthesis | 2 |
| Fatty acid degradation | 2 |
| Arginine and proline metabolism | 2 |
| Glutathione metabolism | 2 |
| Glycosylphosphatidylinositol (GPI)-anchor biosynthesis | 2 |
| Spliceosome | 2 |
| Mismatch repair | 2 |
| Homologous recombination | 2 |
| Two-component system | 2 |
| AMPK signaling pathway | 2 |
| Endocytosis | 2 |
| Peroxisome | 2 |
| Ferroptosis | 2 |
| Glucagon signaling pathway | 2 |
| Pathways in cancer | 2 |
| Viral carcinogenesis | 2 |
| Human T-cell leukemia virus 1 infection | 2 |
| Epstein-Barr virus infection | 2 |

**Supplementary Table S13. The biological process GO categories that were enriched in the genes with positive selection in *Aloe vera* (Only statistically significant GO terms p<0.05 are mentioned)**

| GO Term ID | Description | p-value |
|---|---|---|
| GO:0051128 | regulation of cellular component organization | 0.003 |
| GO:0009415 | response to water | 0.004 |
| GO:0032504 | multicellular organism reproduction | 0.007 |
| GO:0051172 | negative regulation of nitrogen compound metabolic process | 0.008 |
| GO:0015748 | organophosphate ester transport | 0.008 |
| GO:0009890 | negative regulation of biosynthetic process | 0.010 |
| GO:0040008 | regulation of growth | 0.011 |
| GO:0016052 | carbohydrate catabolic process | 0.014 |
| GO:0104004 | cellular response to environmental stimulus | 0.016 |
| GO:0071496 | cellular response to external stimulus | 0.024 |
| GO:2000241 | regulation of reproductive process | 0.024 |
| GO:0015979 | photosynthesis | 0.025 |
| GO:1905392 | plant organ morphogenesis | 0.031 |
| GO:0051726 | regulation of cell cycle | 0.031 |
| GO:0006974 | cellular response to DNA damage stimulus | 0.031 |
| GO:0009735 | response to cytokinin | 0.036 |
| GO:0080134 | regulation of response to stress | 0.040 |
| GO:0009409 | response to cold | 0.043 |

| | | |
|---|---|---|
| GO:0009308 | amine metabolic process | 0.044 |
| GO:0006325 | chromatin organization | 0.046 |
| GO:0010038 | response to metal ion | 0.048 |
| GO:0045165 | cell fate commitment | 0.049 |

**Supplementary Table S14. The cellular component GO categories that were enriched in the positively selected genes in *Aloe vera* (Only statistically significant GO terms p<0.05 are mentioned)**

| GO Term ID | Description | p-value |
|---|---|---|
| GO:0030133 | transport vesicle | 0.031 |
| GO:0005635 | nuclear envelope | 0.046 |

**Supplementary Table S15. The molecular function GO categories that were enriched in the positively selected genes in *Aloe vera* (Only statistically significant GO terms p<0.05 are mentioned)**

| GO Term ID | Description | p-value |
|---|---|---|
| GO:0005509 | calcium ion binding | 0.006 |
| GO:0016798 | hydrolase activity, acting on glycosyl bonds | 0.037 |
| GO:0017056 | structural constituent of nuclear pore | 0.038 |

**Supplementary Table S16. The distribution of genes containing positively selected codon sites in different eggNOG categories in *Aloe vera***

| eggNOG category | Number of genes |
|---|---|
| Function unknown | 443 |
| Signal transduction mechanisms | 167 |
| Posttranslational modification, protein turnover, chaperones | 157 |
| Transcription | 128 |
| Carbohydrate transport and metabolism | 123 |
| RNA processing and modification | 87 |
| Intracellular trafficking, secretion, and vesicular transport | 81 |
| Amino acid transport and metabolism | 80 |
| Translation, ribosomal structure and biogenesis | 79 |
| Secondary metabolites biosynthesis, transport and catabolism | 65 |
| Lipid transport and metabolism | 63 |
| Energy production and conversion | 59 |
| Inorganic ion transport and metabolism | 48 |
| Replication, recombination and repair | 42 |
| Cell cycle control, cell division, chromosome | 30 |

| | |
|---|---|
| partitioning | |
| Cell wall/membrane/envelope biogenesis | 28 |
| Cytoskeleton | 27 |
| Coenzyme transport and metabolism | 25 |
| Chromatin structure and dynamics | 24 |
| Nucleotide transport and metabolism | 23 |
| Defence mechanisms | 15 |
| Nuclear structure | 1 |

**Supplementary Table S17. The distribution of genes containing positively selected codon sites in different KEGG pathways in *Aloe vera* (Only pathways with more than ten genes are mentioned)**

| KEGG Pathway | Number of Genes |
|---|---|
| Ribosome | 25 |
| Protein processing in endoplasmic reticulum | 24 |
| Purine metabolism | 21 |
| Starch and sucrose metabolism | 18 |
| Spliceosome | 18 |
| RNA transport | 18 |
| Plant hormone signal transduction | 18 |
| Cysteine and methionine metabolism | 17 |
| Amino sugar and nucleotide sugar metabolism | 16 |
| Cell cycle | 16 |
| Glycolysis / Gluconeogenesis | 15 |
| Endocytosis | 15 |
| Thermogenesis | 15 |
| Pathways in cancer | 15 |
| Glycerophospholipid metabolism | 14 |
| Lysosome | 13 |
| Epstein-Barr virus infection | 13 |
| Ubiquitin mediated proteolysis | 12 |
| AMPK signaling pathway | 12 |
| Cell cycle - yeast | 12 |
| Plant-pathogen interaction | 12 |
| Viral carcinogenesis | 12 |
| Pyruvate metabolism | 11 |
| Glycerolipid metabolism | 11 |
| Aminoacyl-tRNA biosynthesis | 11 |
| Ribosome biogenesis in eukaryotes | 11 |
| RNA degradation | 11 |
| MAPK signaling pathway - plant | 11 |
| Meiosis - yeast | 11 |
| Glucagon signaling pathway | 11 |
| Salmonella infection | 11 |

| | |
|---|---|
| Human papillomavirus infection | 11 |
| Alanine, aspartate and glutamate metabolism | 10 |
| mRNA surveillance pathway | 10 |
| PI3K-Akt signaling pathway | 10 |
| Alzheimer disease | 10 |
| Huntington disease | 10 |
| Human immunodeficiency virus 1 infection | 10 |

**Supplementary Table S18. The biological process GO categories that were enriched in the genes containing positively selected codon sites in *Aloe vera* (Only statistically significant GO terms p<0.05 are mentioned)**

| GO Term ID | Description | p-value |
|---|---|---|
| GO:0010038 | response to metal ion | 0.002 |
| GO:0044087 | regulation of cellular component biogenesis | 0.005 |
| GO:0021700 | developmental maturation | 0.005 |
| GO:0015850 | organic hydroxy compound transport | 0.009 |
| GO:0072330 | monocarboxylic acid biosynthetic process | 0.019 |
| GO:0006325 | chromatin organization | 0.019 |
| GO:0006928 | movement of cell or subcellular component | 0.019 |
| GO:0006366 | transcription by RNA polymerase II | 0.022 |
| GO:0071669 | plant-type cell wall organization or biogenesis | 0.027 |
| GO:0044419 | interspecies interaction between organisms | 0.029 |
| GO:1905392 | plant organ morphogenesis | 0.030 |
| GO:0006638 | neutral lipid metabolic process | 0.042 |
| GO:0019932 | second-messenger-mediated signaling | 0.045 |
| GO:0006857 | oligopeptide transport | 0.045 |
| GO:0009812 | flavonoid metabolic process | 0.049 |

**Supplementary Table S19. The cellular component GO categories that were enriched in the genes containing positively selected codon sites in *Aloe vera* (Only statistically significant GO terms p<0.05 are mentioned)**

| GO Term ID | Description | p-value |
|---|---|---|
| GO:0009505 | plant-type cell wall | 0.006 |

**Supplementary Table S20. The molecular function GO categories that were enriched in the genes containing positively selected codon sites in *Aloe vera* (Only statistically significant GO terms p<0.05 are mentioned)**

| GO Term ID | Description | p-value |
|---|---|---|
| GO:0016798 | hydrolase activity, acting on glycosyl bonds | 0.006 |
| GO:0001098 | basal transcription machinery binding | 0.021 |
| GO:0005509 | calcium ion binding | 0.035 |

**Supplementary Table S21. The distribution of genes containing unique substitutions with functional impact in different eggNOG categories in *Aloe vera***

| eggNOG category | Number of genes |
|---|---|
| Function unknown | 642 |
| Signal transduction mechanisms | 242 |
| Posttranslational modification, protein turnover, chaperones | 224 |
| Carbohydrate transport and metabolism | 169 |
| Translation, ribosomal structure and biogenesis | 143 |
| Transcription | 142 |
| RNA processing and modification | 131 |
| Intracellular trafficking, secretion, and vesicular transport | 122 |
| Amino acid transport and metabolism | 111 |
| Lipid transport and metabolism | 97 |
| Energy production and conversion | 85 |
| Inorganic ion transport and metabolism | 80 |
| Secondary metabolites biosynthesis, transport and catabolism | 75 |
| Replication, recombination and repair | 68 |
| Cell cycle control, cell division, chromosome partitioning | 60 |
| Coenzyme transport and metabolism | 43 |
| Cytoskeleton | 41 |
| Nucleotide transport and metabolism | 37 |
| Chromatin structure and dynamics | 36 |
| Cell wall/membrane/envelope biogenesis | 32 |
| Defence mechanisms | 18 |

**Supplementary Table S22. The distribution of genes containing unique substitutions with functional impact in different KEGG pathways in *Aloe vera* (Only pathways with more than ten genes are mentioned)**

| KEGG Pathway | Number of Genes |
|---|---|
| RNA transport | 36 |
| Spliceosome | 31 |
| Protein processing in endoplasmic reticulum | 30 |
| Ribosome | 29 |
| Purine metabolism | 26 |
| Ribosome biogenesis in eukaryotes | 24 |
| Alzheimer disease | 24 |
| Glycolysis / Gluconeogenesis | 23 |
| Cysteine and methionine metabolism | 23 |
| Starch and sucrose metabolism | 22 |
| Ubiquitin mediated proteolysis | 22 |

| | |
|---|---|
| Huntington disease | 22 |
| mRNA surveillance pathway | 21 |
| Endocytosis | 20 |
| Cell cycle | 20 |
| Cell cycle - yeast | 20 |
| Pyruvate metabolism | 19 |
| Autophagy - yeast | 19 |
| Thermogenesis | 19 |
| Amino sugar and nucleotide sugar metabolism | 18 |
| Glycerolipid metabolism | 18 |
| Aminoacyl-tRNA biosynthesis | 18 |
| Lysosome | 17 |
| RNA degradation | 16 |
| Plant hormone signal transduction | 16 |
| Peroxisome | 16 |
| Meiosis - yeast | 16 |
| Human T-cell leukemia virus 1 infection | 16 |
| Human papillomavirus infection | 16 |
| Viral carcinogenesis | 15 |
| Salmonella infection | 15 |
| Epstein-Barr virus infection | 15 |
| Carbon fixation in photosynthetic organisms | 14 |
| Glycerophospholipid metabolism | 14 |
| Porphyrin and chlorophyll metabolism | 14 |
| AMPK signaling pathway | 14 |
| Human immunodeficiency virus 1 infection | 14 |
| Alanine, aspartate and glutamate metabolism | 13 |
| Glycine, serine and threonine metabolism | 13 |
| Phenylalanine, tyrosine and tryptophan biosynthesis | 13 |
| Glutathione metabolism | 13 |
| MAPK signaling pathway - plant | 13 |
| HIF-1 signaling pathway | 13 |
| Glucagon signaling pathway | 13 |
| Plant-pathogen interaction | 13 |
| Pathways in cancer | 13 |
| DNA replication | 12 |
| Nucleotide excision repair | 12 |
| Cellular senescence | 12 |
| Insulin signaling pathway | 12 |
| Shigellosis | 12 |
| Glyoxylate and dicarboxylate metabolism | 11 |
| Oxidative phosphorylation | 11 |
| Valine, leucine and isoleucine degradation | 11 |
| Arginine biosynthesis | 11 |

| | |
|---|---|
| Terpenoid backbone biosynthesis | 11 |
| Proteasome | 11 |
| Homologous recombination | 11 |
| mTOR signaling pathway | 11 |
| Autophagy - animal | 11 |
| Oocyte meiosis | 11 |
| Circadian rhythm - plant | 11 |
| Pentose phosphate pathway | 10 |
| Fructose and mannose metabolism | 10 |
| Inositol phosphate metabolism | 10 |
| Methane metabolism | 10 |
| N-Glycan biosynthesis | 10 |
| Various types of N-glycan biosynthesis | 10 |
| FoxO signaling pathway | 10 |
| Longevity regulating pathway - worm | 10 |

**Supplementary Table S23. The biological process GO categories that were enriched in the genes containing unique substitutions with functional impact in *Aloe vera* (Only statistically significant GO terms p<0.05 are mentioned)**

| GO Term ID | Description | p-value |
|---|---|---|
| GO:0009624 | response to nematode | 0.001 |
| GO:0030001 | metal ion transport | 0.003 |
| GO:0016051 | carbohydrate biosynthetic process | 0.004 |
| GO:0015849 | organic acid transport | 0.006 |
| GO:0051187 | cofactor catabolic process | 0.007 |
| GO:0009617 | response to bacterium | 0.008 |
| GO:0051128 | regulation of cellular component organization | 0.011 |
| GO:0072521 | purine-containing compound metabolic process | 0.012 |
| GO:0046777 | protein autophosphorylation | 0.016 |
| GO:0051094 | positive regulation of developmental process | 0.018 |
| GO:0070085 | Glycosylation | 0.022 |
| GO:0007166 | cell surface receptor signaling pathway | 0.026 |
| GO:0042180 | cellular ketone metabolic process | 0.026 |
| GO:0006820 | anion transport | 0.026 |
| GO:0048878 | chemical homeostasis | 0.028 |
| GO:0009308 | amine metabolic process | 0.028 |
| GO:0005976 | polysaccharide metabolic process | 0.030 |
| GO:0009100 | glycoprotein metabolic process | 0.033 |
| GO:0051240 | positive regulation of multicellular organismal process | 0.033 |
| GO:0006090 | pyruvate metabolic process | 0.034 |
| GO:0071669 | plant-type cell wall organization or biogenesis | 0.034 |
| GO:0062012 | regulation of small molecule metabolic process | 0.034 |
| GO:0044262 | cellular carbohydrate metabolic process | 0.036 |

| | | |
|---|---|---|
| GO:0016049 | cell growth | 0.038 |
| GO:0010038 | response to metal ion | 0.040 |
| GO:0006631 | fatty acid metabolic process | 0.043 |
| GO:0022603 | regulation of anatomical structure morphogenesis | 0.048 |

**Supplementary Table S24. The cellular component GO categories that were enriched in the genes containing unique substitutions with functional impact in *Aloe vera* (Only statistically significant GO terms p<0.05 are mentioned)**

| GO Term ID | Description | p-value |
|---|---|---|
| GO:0005802 | trans-Golgi network | 0.002 |
| GO:0000325 | plant-type vacuole | 0.005 |
| GO:0005768 | Endosome | 0.018 |
| GO:0098552 | side of membrane | 0.021 |
| GO:0000139 | Golgi membrane | 0.040 |
| GO:0099023 | tethering complex | 0.042 |

**Supplementary Table S25. The molecular function GO categories that were enriched in the genes containing unique substitutions with functional impact in *Aloe vera* (Only statistically significant GO terms p<0.05 are mentioned)**

| GO Term ID | Description | p-value |
|---|---|---|
| GO:0008194 | UDP-glycosyltransferase activity | 0.002 |
| GO:0022803 | passive transmembrane transporter activity | 0.012 |
| GO:0042562 | hormone binding | 0.017 |
| GO:0043177 | organic acid binding | 0.018 |
| GO:0052689 | carboxylic ester hydrolase activity | 0.020 |
| GO:0005102 | signaling receptor binding | 0.022 |
| GO:0016874 | ligase activity | 0.022 |
| GO:0030551 | cyclic nucleotide binding | 0.034 |
| GO:0043178 | alcohol binding | 0.034 |
| GO:0016758 | transferase activity, transferring hexosyl groups | 0.035 |

**Supplementary Table S26. The distribution of MSA genes in different eggNOG categories in *Aloe vera***

| eggNOG category | Number of genes |
|---|---|
| Function unknown | 31 |
| Translation, ribosomal structure and biogenesis | 13 |
| Signal transduction mechanisms | 11 |
| Carbohydrate transport and metabolism | 11 |
| RNA processing and modification | 11 |
| Posttranslational modification, protein turnover, chaperones | 10 |
| Inorganic ion transport and metabolism | 9 |

| | |
|---|---|
| Transcription | 6 |
| Intracellular trafficking, secretion, and vesicular transport | 6 |
| Cytoskeleton | 5 |
| Coenzyme transport and metabolism | 5 |
| Energy production and conversion | 5 |
| Amino acid transport and metabolism | 4 |
| Replication, recombination and repair | 4 |
| Cell cycle control, cell division, chromosome partitioning | 3 |
| Lipid transport and metabolism | 3 |
| Cell wall/membrane/envelope biogenesis | 3 |
| Secondary metabolites biosynthesis, transport and catabolism | 2 |
| Nucleotide transport and metabolism | 1 |
| Chromatin structure and dynamics | 1 |

**Supplementary Table S27. The distribution of MSA genes in different KEGG pathways in *Aloe vera* (Only pathways with more than one gene are mentioned)**

| KEGG Pathway | Number of genes |
|---|---|
| Ribosome | 7 |
| RNA transport | 5 |
| Cysteine and methionine metabolism | 4 |
| Glycolysis / Gluconeogenesis | 3 |
| Spliceosome | 3 |
| Aminoacyl-tRNA biosynthesis | 3 |
| Circadian rhythm - plant | 3 |
| Plant-pathogen interaction | 3 |
| Human papillomavirus infection | 3 |
| Fructose and mannose metabolism | 2 |
| Starch and sucrose metabolism | 2 |
| Methane metabolism | 2 |
| Glutathione metabolism | 2 |
| Two-component system | 2 |
| Cell cycle | 2 |
| Cellular senescence | 2 |
| GABAergic synapse | 2 |
| Thermogenesis | 2 |
| Central carbon metabolism in cancer | 2 |
| Alzheimer disease | 2 |
| Huntington disease | 2 |
| Pathogenic Escherichia coli infection | 2 |
| Salmonella infection | 2 |

**Supplementary Table S28. The biological process GO categories that were enriched in the MSA genes in *Aloe vera* (Only statistically significant GO terms p<0.05 are mentioned)**

| GO term ID | Description | p-value |
|---|---|---|
| GO:0015748 | organophosphate ester transport | 0.00263 |
| GO:0010038 | response to metal ion | 0.00621 |
| GO:0032504 | multicellular organism reproduction | 0.00868 |
| GO:0046700 | heterocycle catabolic process | 0.01165 |
| GO:0048589 | developmental growth | 0.01165 |
| GO:0019439 | aromatic compound catabolic process | 0.01252 |
| GO:0044270 | cellular nitrogen compound catabolic process | 0.01252 |
| GO:0016052 | carbohydrate catabolic process | 0.01535 |
| GO:0016049 | cell growth | 0.01538 |
| GO:1901361 | organic cyclic compound catabolic process | 0.01643 |
| GO:0009624 | response to nematode | 0.01748 |
| GO:0045165 | cell fate commitment | 0.02348 |
| GO:2000241 | regulation of reproductive process | 0.02462 |
| GO:0080134 | regulation of response to stress | 0.02652 |
| GO:0042157 | lipoprotein metabolic process | 0.02825 |
| GO:0007017 | microtubule-based process | 0.02968 |
| GO:0008283 | cell proliferation | 0.02968 |
| GO:0072524 | pyridine-containing compound metabolic process | 0.02968 |
| GO:1901698 | response to nitrogen compound | 0.03029 |
| GO:0070482 | response to oxygen levels | 0.03337 |
| GO:0009735 | response to cytokinin | 0.03447 |
| GO:0015931 | nucleobase-containing compound transport | 0.03869 |
| GO:0090351 | seedling development | 0.03869 |
| GO:0022603 | regulation of anatomical structure morphogenesis | 0.03883 |
| GO:0021700 | developmental maturation | 0.04894 |

**Supplementary Table S29. The molecular function GO categories that were enriched in the MSA genes in *Aloe vera* (Only statistically significant GO terms p<0.05 are mentioned)**

| GO term ID | Description | p-value |
|---|---|---|
| GO:0015605 | organophosphate ester transmembrane transporter activity | 0.01886 |
| GO:0005200 | structural constituent of cytoskeleton | 0.02383 |
| GO:0015932 | nucleobase-containing compound transmembrane transporter activity | 0.04144 |
| GO:1901505 | carbohydrate derivative transmembrane transporter activity | 0.04812 |

**Supplementary Table S30. The number of coding genes in monocots including *Aloe vera*. The coding genes for the other monocots were identified using Ensembl plants database [5].**

| Species | Number of coding genes |
|---|---|
| *Aloe vera* | 86,177 |
| *Dioscorea rotundata* | 19,086 |
| *Arabidopsis thaliana* | 27,655 |
| *Leersia perriei* | 29,078 |
| *Aegilops tauschii* | 29,630 |
| *Panicum halii HAL2* | 32,263 |
| *Panicum halii FIL2* | 33,805 |
| *Sorghum bicolor* | 34,118 |
| *Brychopodium distychyon* | 34,310 |
| *Setaria italicaa* | 35,831 |
| *Musa acuminata* | 36,525 |
| *Oryza sativa japonica* | 37,960 |
| *zea mays* | 39,591 |
| *Hordeum vulgare* | 39,841 |
| *Oryza sativa indica* | 40,745 |
| *Ergarostis tef* | 41,935 |
| *Saccharum spontaneum* | 83,815 |
| *Triticum aestivum* | 107,891 |

**SUPPLEMENTARY FIGURES**

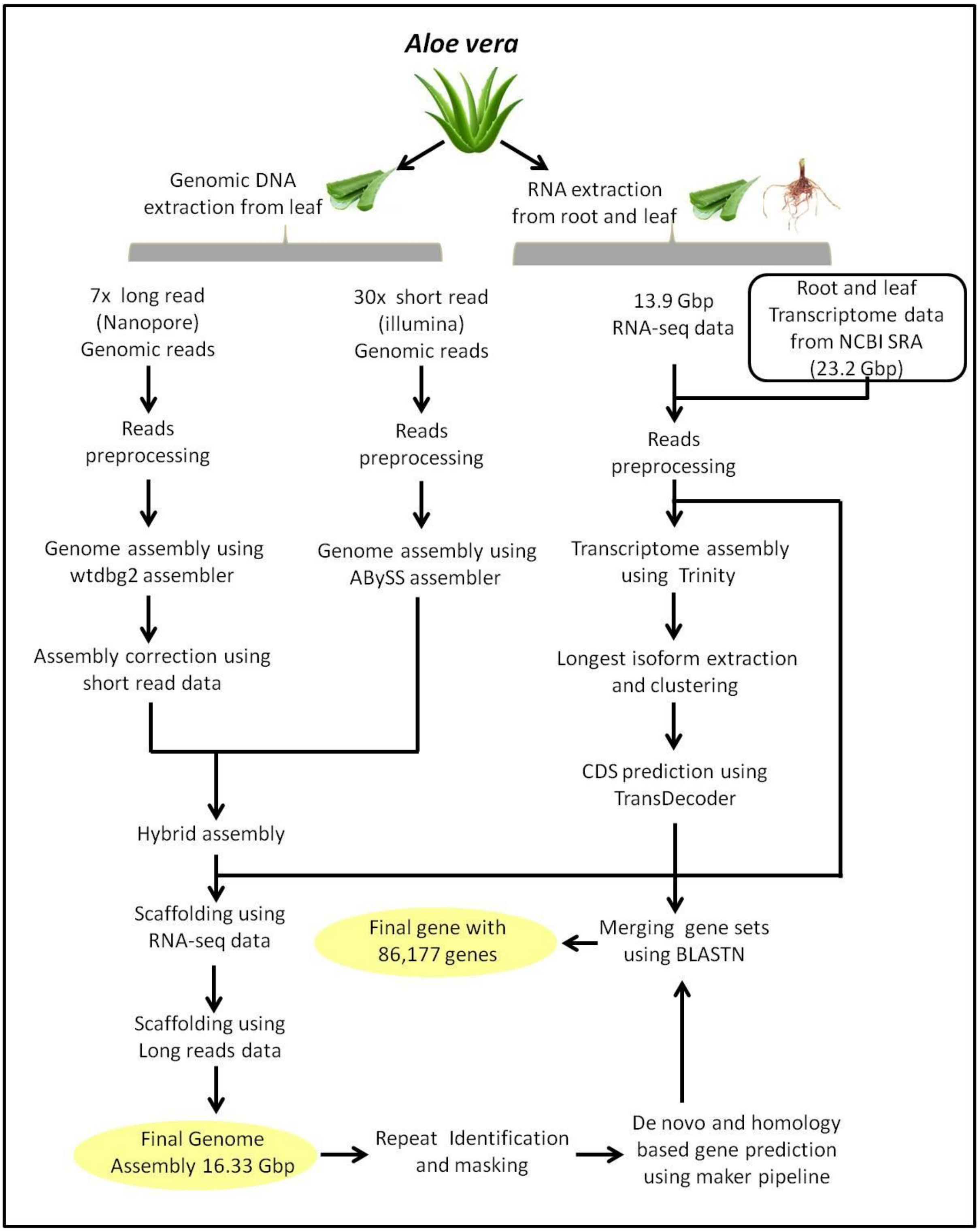


**Supplementary Figure S1. The complete workflow of the genomic and transcriptomic data analysis for *Aloe vera***

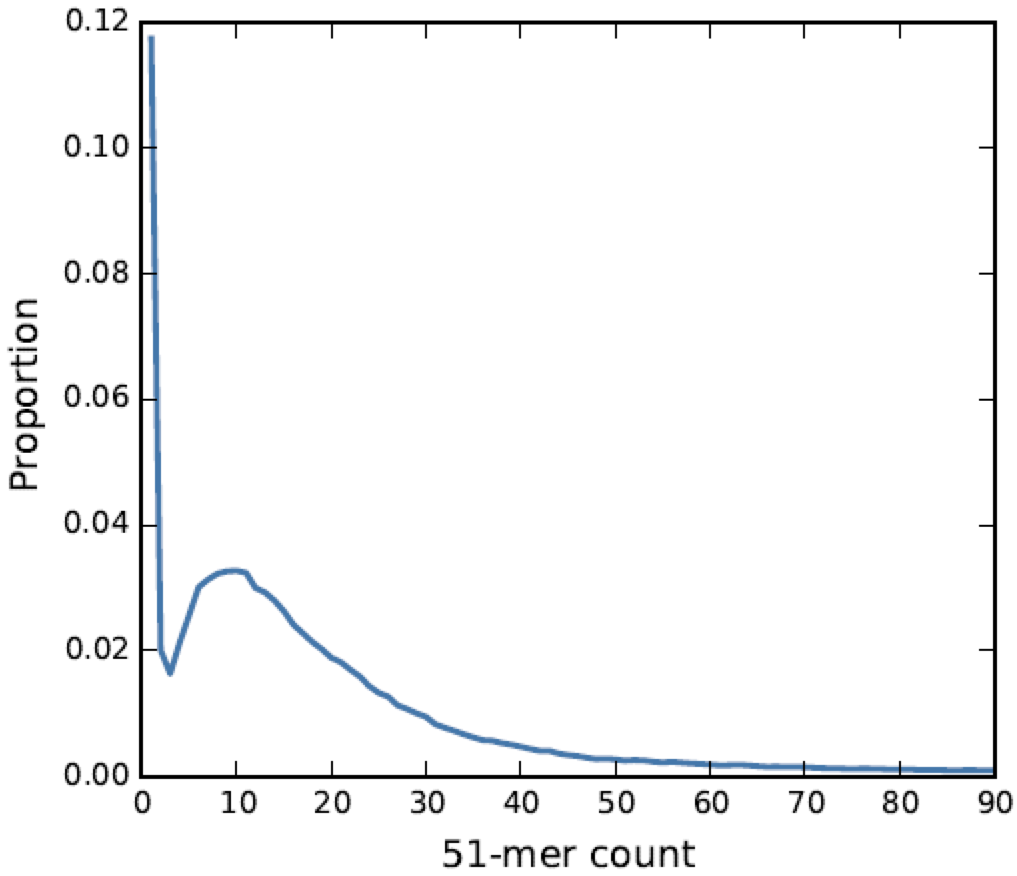


**Supplementary Figure S2. k-mer count distribution for the 51-mer. The y-axis is the proportion of 51-mers and x-axis is the count of the 51-mer.**

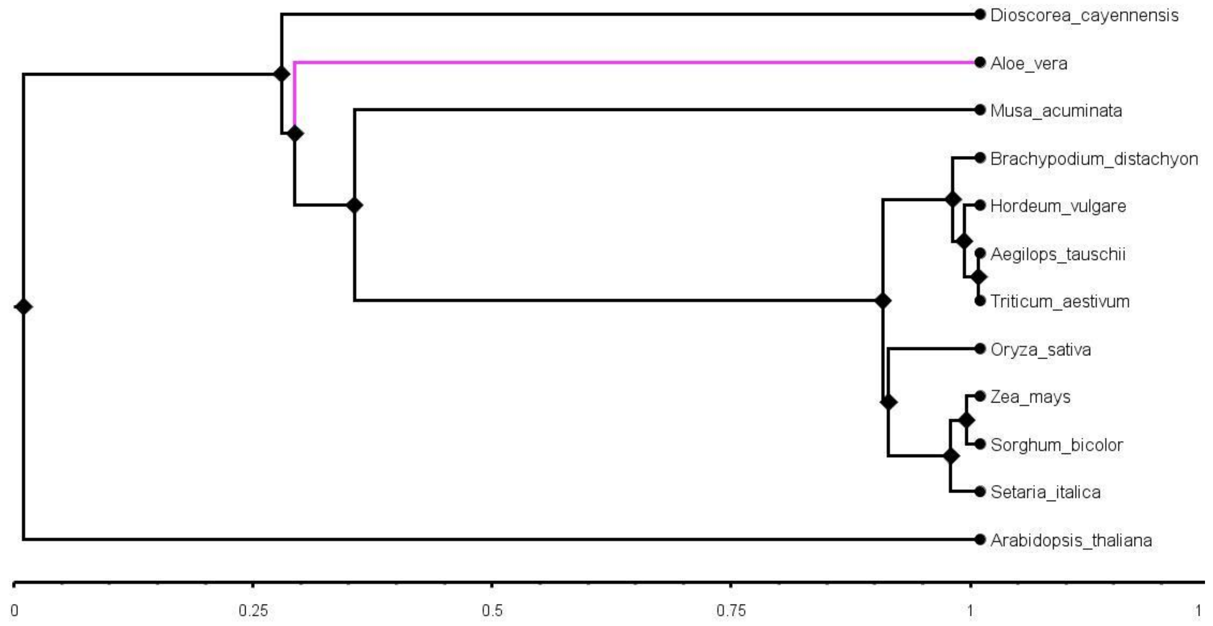


**Supplementary Figure S3. The phylogenetic tree of the monocot species that were common in our study and plant megaphylogeny is shown [6]. The *Arabidopsis thaliana* was used as an outgroup.**

**SUPPLEMENTARY TEXT**

**Supplementary Text S1.**

**Sample collection and Species identification**

The plant was bought from a local nursery in Bhopal, India. The plant was kept in the lab for DNA and RNA extraction. The DNA was initially isolated from DNeasy Plant mini kit (Qiagen, United States). The pulp or gel from the leaf was scrapped out before grinding. While grinding with liq. Nitrogen, 1 ml of AP1 buffer was added to increase the yield. Further steps were followed as given in the kit. The DNA was eluted in 50 µl elution buffer (Qiagen, United States). The DNA was quantified on Qubit 2.0 fluorometer by using Qubit DNA BR assay kit (Life Technologies, United States). The DNA was used for amplification of complete ITS1 and ITS2 (Internal Transcribed Spacer) and Maturase K (MatK) regions by using following primer sets:

| | | |
|---|---|---|
| (i) | Complete ITS region: | forward primer 5′-TCCGTAGGTGAACCTGCGG-3′ |
| | | reverse primer 5′-TCCTCCGCTTATTGATATGC-3′ |
| (ii) | ITS1 region: | forward primer 5′-TCCGTAGGTGAACCTGCGG-3′ |
| | | reverse primer 5′-GCTGCGTTCTTCATCGATGC-3′ |
| (iii) | ITS2 region: | forward primer 5′-GCATCGATGAAGAACGCAGC-3′ |
| | | reverse primer 5′-TCCTCCGCTTATTGATATGC-3′ |
| (iv) | Mat K region: | forward primer 5′-CGATCTATTCATTCAATATTTC-3′ |
| | | reverse primer 5′-TCTAGCACACGAAAGTCGAAGT-3′ |

The PCR programme run on Veriti 96 well thermal cycler (Applied Biosystems) for ITS regions was 94 ˚C for 3 mins, 35 cycles of 94 ˚C for 1 min, 55 ˚C for 1 min and 72 ˚C for 2.5 mins and 72 ˚C for 10 mins. Similarly, the programme for MatK was 95 ˚C for 3 mins, 35 cycles of 95 ˚C for 30 sec, 50 ˚C for 3 mins and 72 ˚C for 1:15 min and final extension at 72 ˚C for 7 mins. The amplification products were assessed by running them on 2% agarose gel electrophoresis. The amplified products were purified and sequenced at in-house Sanger sequencing facility. All the sequences were checked for alignment with NCBI database using BLASTN and showed highest identity with *Aloe vera* which confirmed the species as *Aloe vera*.

**Genome sequencing**

*Short read sequencing*

DNA extraction: The DNA was initially isolated from DNeasy Plant mini kit (Qiagen, United States). The pulp or gel from the leaf was scrapped out before grinding. While grinding with liq. Nitrogen, 1 ml of AP1 buffer was added to increase the yield. Further steps were followed as given in the kit. The DNA was eluted in 50 µl elution buffer (Qiagen, United States). The DNA was quantified using Qubit DNA BR assay kit on Qubit 2.0 fluorometer (Life Technologies, United States). The library was prepared using NEBNext Ultra II DNA Library preparation Kit for Illumina (New England Biolabs, England) and TruSeq DNA Nano Library preparation kits (Illumina, Inc., United States). The library size was evaluated by Agilent 2100 Bioanalyzer and library was quantified by qPCR. The library was sequenced on Illumina HiSeq X ten platform and NovaSeq 6000 (Illumina, Inc., United States) for 150 bp paired end reads.

*Long read sequencing*

The DNA extraction for long read sequencing was done by using Carlson lysis buffer [ 100 mM Tris; 2% CTAB; 1.4 M NaCl; 1% PEG 8000; 20 mM Ethylene read sequencing Diamine Tetra Acetic acid (EDTA)]. To 50 ml of Carlson buffer, β-mercaptoethanol (125 µl) was added and vortexed to mix it properly. The plant part taken for DNA extraction was leaf. The pulp or gel from the leaf was scrapped out before homogenization. The sample was homogenized in liquid nitrogen by using a mortar and pestle (autoclaved and precooled at -20˚C for 30 mins) and transferred to a microcentrifuge tube. Carlson lysis buffer (1 ml) was preheated at 65˚C for 30 mins and added to the sample. After adding 2 µl of RNase A (20 mg/ml) and 25 ul of Proteinase K (20 µL/mL) and vortexing for 5 sec, the sample was incubated at 65˚C for 1 hr. The sample was mixed in between by inverting 10 times. After incubation, the sample was allowed to cool at room temperature for 5 mins. The sample tube was provided with 1 ml of chloroform, vortexed and centrifuged at 5,000 xg for 15 mins at 4˚C. The top aqueous layer was transferred with wide bore tip to a new centrifuge tube. The 0.7X volume of isopropanol was added, mixed by inverting 10 times and incubated at -20˚C for overnight. The tube was centrifuged at 5,000 xg for 45 mins at 4˚C. In conventional method, the supernatant was discarded and pellet was washed with 1 ml of ice-cold 70 % ethanol by centrifuging at 5,000 xg for 10 mins at 4˚C. The supernatant was again discarded and the pellet was air dried to evaporate all of the ethanol residues. The DNA was eluted in 50 µl of nuclease free water.

In kit-based method, Blood and Cell culture kit with Genomic tip 20 (Qiagen, United States) was used. The Pelleted DNA was not washed with 70% ethanol but it was dissolved in G2 buffer by incubating at 50˚C for 30 mins. The dissolved DNA was passed through equilibrated Genomic tip 20. The column was washed thrice with QC buffer (1 ml) and eluted in 1 ml of QF buffer (pre heated at 56˚C). The DNA was allowed to precipitate in 0.7X Isopropanol for overnight at -20˚C. The precipitated DNA was washed and eluted same as in conventional method.

The DNA was quantified with Qubit 2.0 fluorometer by using Qubit DNA BR assay kit (Life Technologies, United States). The DNA quality was checked by running on agarose gel and NanoDrop™ 8000 Spectrophotometer (ThermoFisher Scientific, USA). To reach the required purity of samples, they were purified with Ampure XP beads (Beckman Coulter, USA). The purified samples were used for library preparation by following the protocol Genomic DNA by Ligation using SQK-LSK109 kit (Oxford Nanopore). The library was loaded on FLO-MIN106 Flow cell (R 9.4.1) and sequenced on MinION (Oxford Nanopore, UK) using MinKNOW software (versions 3.4.5 and 3.6.0).

**Transcriptome sequencing**

The leaf and root part of plant were taken as sample for RNA extraction. The samples were grinded in liquid nitrogen with the help of mortar and pestle. The powdered sample (100 mg) was transferred to centrifuge tube to which 1 ml of TRIzol reagent (Invitrogen, USA) was added and shaken for 5 mins. For complete dissociation of nucleoprotein complexes the tubes were incubated for 5 mins at room temperature. Chloroform (200 µl) was added to the tubes and vortexed for 15 sec and incubated at room temperature for 10 mins. After incubation the tubes were centrifuged at 12,000 xg for 15 mins at 4˚C and upper aqueous phase was transferred to a new centrifuge tube. Isopropanol (500 µl) was added, mixed thoroughly and allowed to precipitate for 5-10 mins at room temperature. The RNA was pellet down by centrifuging at 12,000 xg for 10 mins at 4˚C and supernatant was discarded. The pellet was washed with 1ml of 75% ethanol by centrifuging at 7,500 xg for five mins at 4˚C. The supernatant was discarded and was kept at 37˚C for 30 mins to evaporate the residual ethanol. The RNA pellet was resuspended in 30 ul of nuclease free water, dissolved the pellet by pipette mixing and incubated at 55-60˚C for 10-15 mins [7]. The RNA was diluted 10 times and quantified on Qubit 2.0 fluorometer by using Qubit HS Assay kit (Invitrogen, USA). The library was prepared by using TruSeq Stranded mRNA LT Sample Prep kit and following TruSeq Stranded mRNA Sample Preparation Guide (Illumina, Inc., United States) and sequenced on Illumina NovaSeq 6000 platform for 101 basepair paired end reads.

**Supplementary Text S2:**

**Data pre-processing**

The raw Illumina sequence data was processed using the Trimmomatic V0.38 [8]. The adapters used for the sequencing were trimmed using the parameters: 2 mismatches to be allowed in the seed matching with seed length of 16, palindrome clip threshold of 30, and simple clip threshold of 10. The low quality bases or N's were removed from the leading and trailing ends of the reads with the quality threshold of 15. The reads were scanned with a sliding window of 4 bp and the reads were trimmed when average PHRED quality score per base went below 15. After these steps, all the reads smaller than 60 bp were removed.

For nanopore data the raw sequencing reads were obtained in fast5 from the MinKNOW v3.6.0 and basecalling was performed using Guppy v3.2.1. Adapter sequences were then removed based on all known adapters by using Porechop v0.2.3.

**Genome Size Estimation**

The SGA-preqc was used to estimate the genome size of *Aloe vera* species [9]. It uses a k-mer count distribution method, where only the k-mers with higher occurrences are considered for genome size estimation. Thus, it reduces the impact of sequencing errors on the estimation which is very useful for higher repeat containing plant genomes. At first the sga preprocess was run (with the option -pe-mode set to 0 to consider all the paired-end and single-end filter reads for the analysis) to preprocess the raw reads. Next, the sga index was run with 'ropebwt' algorithm and --no-reverse option to index the preprocessed reads, and finally, the sga preqc was run with default options for genome size estimation.

**Genome assembly**

The filtered paired and unpaired Illumina reads were *de novo* assembled using ABySS v2.1.5 with bloom-filter function for a Bloom filter size of 950 GB, the bloom filter hash function and minimum k-mer count threshold for bloom-filter assembly were used as default [10]. The other parameters were: minimum alignment length of a read of 40 bp, minimum unitig size required for building contigs of 500 bp, and minimum contig size required for building scaffolds of 500 bp. Different assemblies were generated on a sample dataset at different k-mer values: 41, 87, 96, 107, 117, 127. The best assembly resulted on k-mer value of 107 hence, the final assembly on complete data was performed at the k-mer value of 107.

The preprocessed nanopore reads were *de novo* assembled using wtdbg2 v2.0.0 with an estimated genome size of 16 GB, subsampling k-mer value of 1.0, minimum read depth of a valid edge of 2, with keeping the contained reads during alignments. The minimum length of alignment between reads was set to 2,048 bp.

To generate the hybrid assembly from the short-read and long-read assembly generated above, the contigs from the ABySS and wtdbg2 were merged using BLASTN. All the ABySS contigs were searched in the wtdbg2 contigs and all the matching contigs with the criteria of 50% query coverage, e-value of $<10^{-6}$ and 90% identity were removed, and the unique contigs from ABySS and wtdbg2 assembly were merged to construct a hybrid assembly.

**Genome polishing**

The obtained genome assembly was first corrected for the assembly and sequencing errors (generally introduced by the assembler, read error correction tools, or the sequencing technology) using short read data. The hybrid assembly was indexed and the filtered short-read data was mapped on to the hybrid assembly using hisat2 v2.1.0 to construct the ".sam" file [11]. The ".sam" was converted to ".bam" using samtools v1.9 [12]. The mapped ".bam" file was used to perform the correction of the long-reads assembly by using SeqBug [13]. The corrected assembly from the long-reads was used to generate the final hybrid assembly using BLASTN. Further, the RNA-seq data was used to perform the scaffolding of the obtained assembly. The complete RNA-seq data from this study and other studies that was used for the transcriptome assembly was mapped on the indexed SeqBug-polished assembly using hisat2 v2.1.0 [3, 4]. The combined ".bam" file was generated using samtools v1.9. The combined ".bam" file was used for the RNA-seq data based scaffolding using 'Rascaf' [14]. To further improve the assembly and to remove the undetermined bases in the assembly, the gap closing of the assembly was performed using the long-read data. The preprocessed nanopore data was used for the long-read based gap-closing to generate the final *Aloe vera* genome assembly using LR_Gapcloser (https://github.com/CAFS-bioinformatics/LR_Gapcloser/).

**Supplementary Text S3:**

**Genome annotation**

*Repeats Identification*

The tandem repeats in the genome were identified using the Tandem Repeat Finder (TRF) v4.09 with the parameters: matching weight = 2, mismatching penalty = 7, indel penalty = 7, match probability = 0.8, indel probability = 0.1, minimum alignment score = 50, and maximum period size = 2,000 [15].

*The Identification of transfer RNAs (tRNAs)*

The tRNAs are very large and complex non-coding RNA families and present in all the living organisms. Thus, tRNAs in the *Aloe vera* genome were predicted on the final gap-closed assembly using tRNAscan-SE v2.0.5 on the default parameters [16, 17]. The tRNAscan-SE uses a companion Genomic tRNA Database and UCSC genome browser to identify the tRNAs present in a genome. In this method a total of 3,119 tRNAs were identified, of which standard amino acids related tRNAs were 1,978, possible suppressor tRNAs (CTA, TTA, TCA) were 9, undetermined/unknown isotypes tRNAs were 29, and predicted pseudogenes tRNAs were 1,103. Further, a total of 128 tRNAs with introns were also identified.

*The identification of small nucleolar RNAs (snoRNAs), Small nuclear RNAs (snRNAs), and signal recognition particle RNAs (srpRNAs)*

The snoRNAs are one of the abundant types of non-coding RNAs that function primarily to direct the modification of other non-coding RNAs specifically the snRNAs, ribosomal RNAs (rRNAs), and tRNAs. The snoRNA sequences were retrieved from the Ensembl plant database for the representative species from all the monocot genera for which the ncRNA data was available [5, 18]. The redundancies were removed by performing the clustering using CD-HIT-EST v4.8.1, and the non-redundant snoRNA sequences were used for the homology-based search against *Aloe vera* genome using BLASTN alignment tool [19, 20]. The snRNAs, srpRNAs, and rRNA were also identified in the *Aloe vera* genome using the similar methodology as used for the identification of snoRNAs.

*The identification of microRNAs (miRNAs)*

A total of 38,589 hairpin miRNAs were retrieved from the miRBase database [21]. These hairpin miRNAs were clustered separately to remove redundancies using CD-HIT-EST v4.8.1 [20]. After clustering, the hairpin miRNAs dataset had a total of 22,365 sequences. The non-redundant sequences were used to identify the hairpin miRNAs in the *Aloe vera* genome using homology-based search by BLASTN alignment tool with the thresholds: identity ≥80% and e-value <1e-03 [22].
